# NLRX1 is an essential, druggable regulator of mitochondrial permeability transition

**DOI:** 10.64898/2026.08.24.746860

**Authors:** Rachel Peltier-Heap, David W. Frederick, Robert J. Pickering, Sonja Ghidelli-Disse, Kirsten Searle, Juliette C. Barber, Nicholas Galwey, Lea P. Wilhelm, Kathy Triantafilou, Martha Triantafilou, Oliver Wright, Sumitra Ramachandran, Yan-Ming Tan, WenHao Xia, Chiu-Cheong Aw, Pamela Oon, Emma Rivers, Kiew Ching Lee, Yannick Lacroix, Esther Reddy, Robert P. Glover, Gino Brunori, Ashley J. Broom, David Grimsditch, Annie Garcia, Avril Robertson, Kazufumi Hirano, Angela A. Brady, Edward Browne, Gerard Drewes, Ian G. Ganley, Kate Schroder, Mahmood Ahmed, Lee M. Booty

**Affiliations:** GSK, Stevenage, SG1 2NY, UK; GSK, Upper Providence, PA, 19426, USA; Institute of Cancer Research, SW7 3RP, London, UK; Cellzome GmbH, a GSK Company, 69117, Heidelberg, Germany; GSK, Biopolis, Singapore; Immunology Network, GSK, Stevenage, SG1 2NY, UK; MRC Protein Phosphorylation and Ubiquitylation Unit, University of Dundee, Dundee, DD1 5EH, UK; School of Medicine, University of Cardiff, CF14 4YS, UK; Trinity College Dublin, Dublin, D02 PN40, Ireland; GSK, Ware, SG12 0DP, UK; School of Chemistry and Molecular Biosciences, The University of Queensland, St Lucia, Queensland, Australia; Institute for Molecular Bioscience and Institute for Molecular Bioscience Centre for Inflammation and Disease Research, The University of Queensland, St. Lucia, Australia; Walter and Eliza Hall Institute, Parkville, Melbourne, Australia

**Keywords:** drug discovery, innate immunity, mPTP, NLRX1, mitochondria

## Abstract

The molecular composition of the mitochondrial permeability transition pore (mPTP) remains contested, and several efficacious mPTP inhibitors act through undefined, cyclophilin D (CypD)-independent targets. Using two structurally distinct chemotypes of optimised, brain-penetrant mPTP inhibitors as chemical probes, we applied affinity-based chemoproteomics to identify the mitochondrial NOD-like receptor NLRX1 as their shared target. Both chemotypes bind NLRX1, and binding potency across a compound series tracks mPTP-inhibitory activity. Using CRISPR-Cas9-edited human cells and Nlrx1-/- mouse tissues, we show that NLRX1 is required for normal calcium-induced mPTP opening: its loss raises the calcium threshold for pore opening and its overexpression lowers it, independently of CypD. NLRX1 associates with postulated mPTP components, including ATP synthase and the adenine nucleotide translocase, in a compound-sensitive manner, and sustains mitochondrial protein homeostasis over longer timescales. The lead compound, GSK900, is orally bioavailable, brain-penetrant, and active in an mPTP-sensitive neurological injury model. These findings, converging with recent genetic studies, establish NLRX1 as an essential, CypD-independent regulator of mitochondrial permeability transition and provide brain-penetrant chemical tools to interrogate this biology.

## Introduction

Mitochondria are key mediators of cellular survival responsible both for the oxidation of organic substrates toward the production of ATP and as central nodes in the execution of cell death pathways, including apoptosis and necrosis ^1^. One route to initiation of necrotic cell death is the process of mitochondrial permeability transition, during which the buildup of cellular stressors, such as reactive oxygen species (ROS) and Ca^2+^, induces the inner mitochondrial membrane to become porous, dissipating the intramembrane electrochemical potential and leading to energetic crisis and eventual cell death ^2,3^. The existence of a highly regulated protein super complex that is critical to this mitochondrial permeability transition, known as the mitochondrial permeability transition pore (mPTP), has been accepted for decades, yet the precise identity of the pore components, as well as their stoichiometry and physical interactions, remain a matter of debate^4^. Nonetheless, it can be deduced from genetic and pharmacological models that the mPTP plays an important role in the pathophysiology of ischemia-reperfusion injury and neurological disease, to name a few ^5–7^.

Specific therapeutic targeting of the mPTP requires a better understanding of the pore components and their intrinsic regulators. Historical attempts to define such constituents of the mPTP have been based on candidate-based approaches, relying on reconstitution of protein complexes which approximate those observed in intact organelles ^8,9^. Among the suggested proteinaceous components of the mPTP are the F_1_F_0_ ATP synthase, voltage-dependent anion channel (VDAC), and adenine nucleotide translocases (ANTs; SLC25A4-6), yet genetic studies have proven each to be non-essential for mPTP formation ^10^. Cyclophilin D (CypD) is the most established positive regulator of the pore and a high-affinity target of the canonical mPTP inhibitor, cyclosporin A (CsA) ^11^. Genetic deletion of CypD or CsA treatment renders mice resistant to a multitude of pathologies, including muscular dystrophy, osteoporosis, and ischemic brain injury ^12–14^. Due to simultaneous off-target inhibition of calcineurin, as well as each of the 15 additional members of the cyclophilin family, CsA yields immunosuppressive and other side effects limiting its therapeutic utility in degenerative disease. Whilst numerous CsA derivatives have been reported, no cyclophilin-D selective, brain penetrant inhibitors have progressed to clinical trial ^15–18^. In recent years, the atypical mitochondrial NOD-like receptor NLRX1 (nucleotide-binding oligomerisation domain-like, leucine-rich-repeat-containing protein X1) has emerged as a candidate regulator of the mPTP. Unusual among NLRs in localising to mitochondria, NLRX1 has been studied principally in innate immune signalling and mitophagy, but two recent and independent lines of work now implicate it directly in permeability transition: genetic studies in the heart show that NLRX1 deletion abolishes calcium-induced mPTP opening^68^, and an unbiased phenotypic CRISPR screen identified NLRX1 as an essential activator of the human permeability transition^69^. These convergent genetic findings motivate an orthogonal, chemical approach to establish and pharmacologically exploit this role.

Given the established pharmacological tractability of the mPTP, unbiased phenotypic screening of small molecule libraries offers the highest probability of discovering selective non-peptide inhibitors. Following this approach, several groups have previously identified structurally distinct classes of small molecules, including cinnamic anilides, isoxazoles, and benzamides, which appear to inhibit mPTP formation or function in isolated mitochondria ^5,19–22^. Compounds from these chemical classes show synergy with CsA, in that there exists an additive inhibitory effect above the maximum inhibition achievable with CsA alone, therefore suggesting that the undefined molecular targets of these inhibitors are independent of CypD. We hypothesised that non-peptide inhibitors pharmacologically optimised for use in neurological disease ^23^ could be employed as chemical probes to reveal novel candidate regulators of mitochondrial calcium homeostasis.

Here we explore two chemically-distinct hits from an mPTP inhibitor screen and, through affinity-based chemoproteomics, identify a high-affinity interaction between these compounds and NLRX1. We highlight a cyclophilin-D independent function of these compounds and provide evidence for a role for NLRX1 in mPTP function, whereby use of these identified compounds or genetic ablation of NLRX1 inhibit murine and human mPTP opening in isolated mitochondria and whole cell systems. Further investigation, including a proteomic interactome of NLRX1 that contains putative mPTP components, uncovered a role for NLRX1 in controlling mitochondrial homeostasis with impact on mitochondrial function, and preliminary evidence that *in vivo* targeting of NLRX1 may be beneficial in neurological injury scenarios.

## Materials and Methods

### Key Resources Table

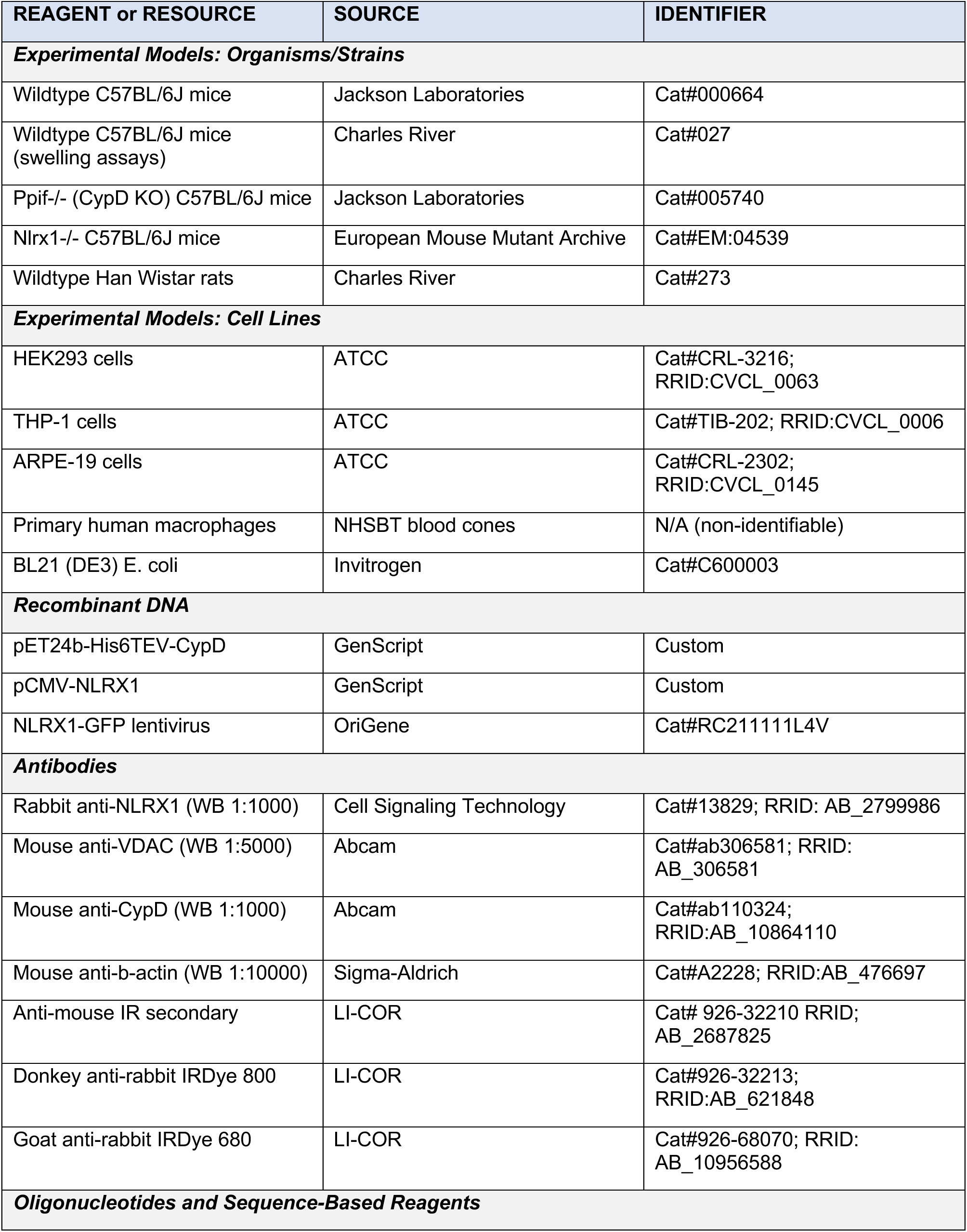
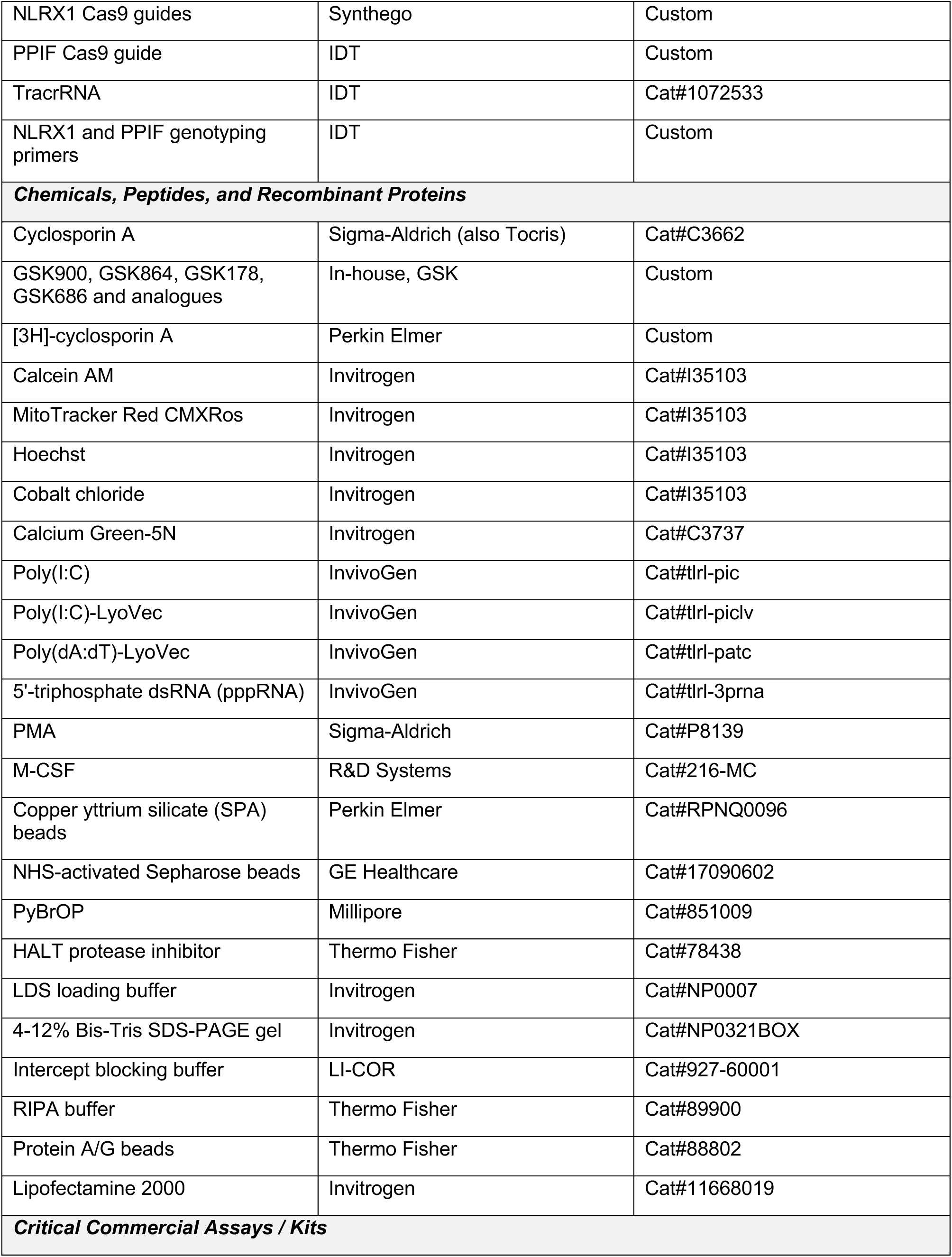
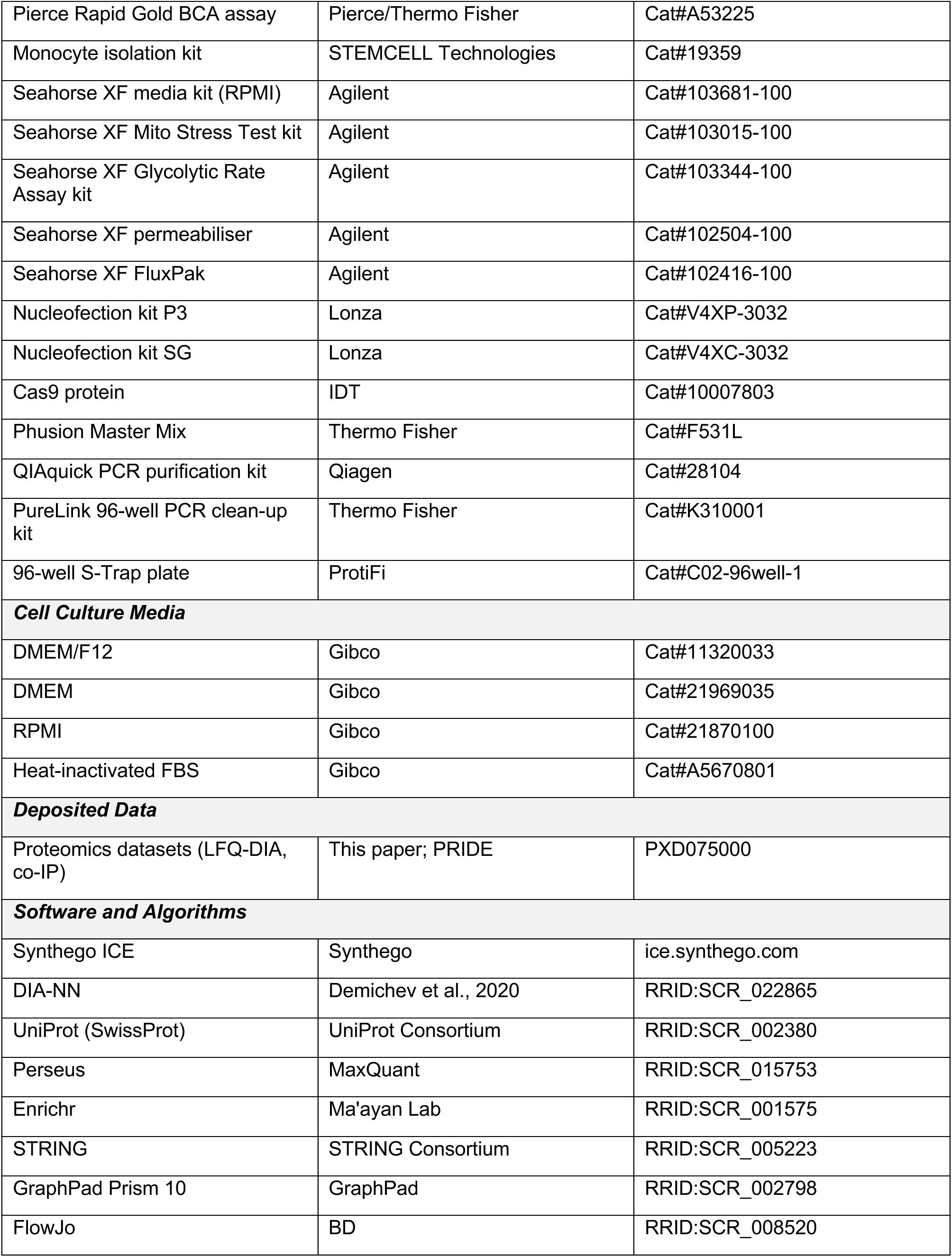
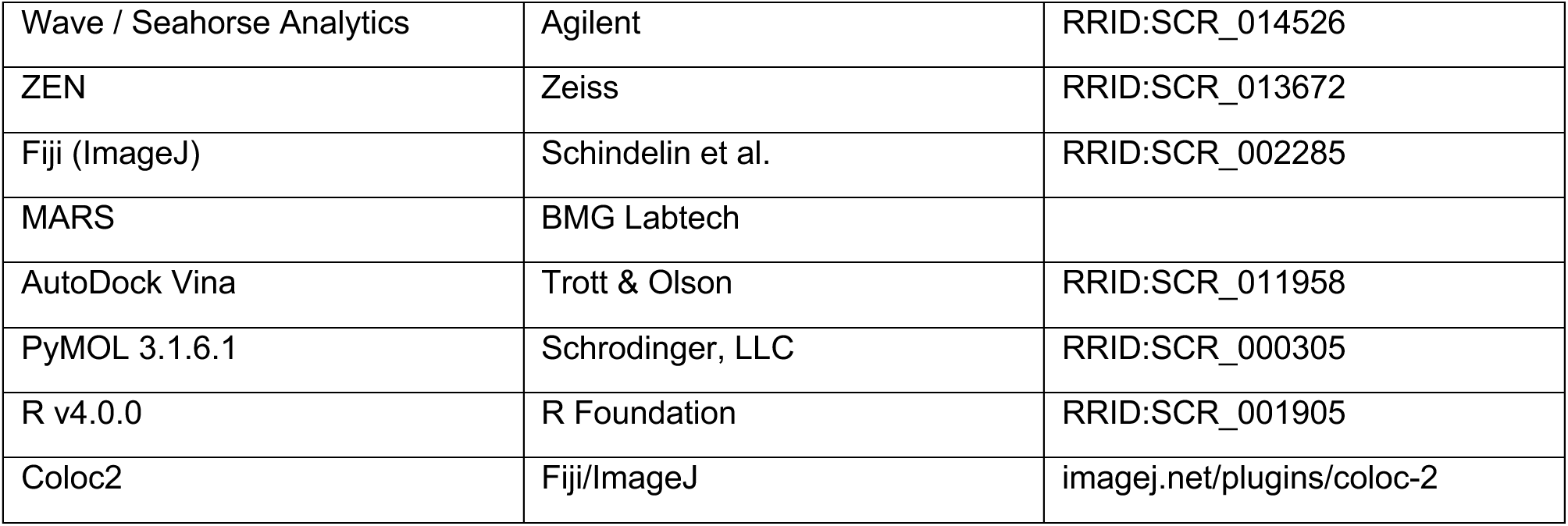

### Resource availability

Further information and requests for resources and reagents should be directed to the Lead Contact, Lee M. Booty.

Proteomics datasets have been deposited at PRIDE and are publicly available as of the date of publication (PXD075000). This paper does not report original code.

### Experimental model details

Mouse strains, rat strains, cell lines (HEK293, THP-1, ARPE-19), and primary human macrophages are as listed in the Key Resources Table; husbandry, culture and derivation details are as described in the existing Materials and Method Details below.

## Materials

All chemicals were purchased from Sigma-Aldrich unless stated here. Cyclosporin A (Tocris); GSK compounds were synthesised in GSK medicinal chemistry laboratories (Singapore); [^3^H]-cyclosporin A (Perkin Elmer); Calcein AM, MitoTracker Red, Hoechst, Cobalt Chloride, Calcium Green-5N (Invitrogen). Mice were obtained from Jackson Laboratories (Wildtype and *Ppif*^-/-^) or the European Mouse Mutant Archive (*Nlrx1*^-/-^), or for swelling assays rats (Han Wistar) and mice (C57BL/6J) were obtained from Charles River. CRISPR guides were obtained from IDT (PPIF) or Synthego (NLRX1), and primers were obtained from IDT. Immune stimuli (Poly(I:C), Poly(I:C)-LyoVec, Poly(dA:dT), 5’-triphosphate dsRNA (pppRNA)) were from Invivogen. Seahorse consumables were from Agilent.

### Animal Use

All animal studies were ethically reviewed and carried out in accordance with the Animals (Scientific Procedures) Act 1986 and the GSK Policy on the Care, Welfare and Treatment of Animals.

### Mitochondrial isolations

Liver (mouse or rat), heart, kidney and brain (all mouse) were isolated after culling by cervical dislocation or decapitation. Tissues were washed in STE buffer (120 mM sucrose, 10 mM Tris, 1 mM EGTA, pH 7.4 (with additional 0.1 % bovine serum albumin (BSA) for the brain and heart)) before chopping. All steps were performed at 4 °C. Tissues were homogenised in STE (except brain, which was homogenised in 12 % Percoll solution) in a Dounce homogeniser until a complete suspension was achieved. Liver and kidney homogenate were centrifuged at 650 x g for 10 min and resulting supernatant centrifuged at 7700 x g for a further 10 min. Pellets were resuspended in STE and centrifuged at 7700 x g for 10 min. Heart homogenate was centrifuged at 700 x g for 5 min and supernatant transferred. Pellet was resuspended in STE and re-centrifuged at 700 x g for 5 min. Supernatants were combined and centrifuged at 5500 x g for 10 min. Brain homogenate was layered on top of a 26:40% Percoll gradient and centrifuged at 21000 x g for 10 min. Mitochondrial fraction was obtained and re-centrifuged at 15000 x g for 10 min. All mitochondrial pellets were resuspended in STE before protein concentration determination by BCA assay.

For mitochondrial isolations from HEK293 cells, cells were cultured in 3 x T500 triple layer flasks before harvesting using TrypLE detachment buffer (Gibco). Cells were pelleted by centrifugation at 300 x g for 5 min and resuspended in 50 mL cold PBS before centrifugation at 300 x g for 5 min. Mitochondrial isolation protocol from cells was followed as outlined in Clayton & Shadel (2014)^24^ and final protein concentration calculated by BCA assay.

### Swelling Assay

Mouse mitochondria were isolated as above and resuspended to 0.5 mg protein/mL in 130 mM sucrose, 50 mM KCl, 2.5 mM KH_2_PO_4_, 5 mM HEPES pH 7.4. Using a FlexStation 3 (Molecular Devices), Swelling Buffer (Buffer as above with 5 µM rotenone, 20 mM sodium succinate and compounds of interest) was added to 50 µg of mitochondria followed by addition of 200 µM CaCl_2_ and 5 mM KH_2_PO_4._ Absorbance at 540 nm was measured throughout additions and data here show post-addition of KH_2_PO_4._

### Calcium Retention Capacity (CRC) Assay

Mitochondria were isolated from mouse tissue or HEK293 cells as above. For additive CRC assays, mitochondria were resuspended in 150 mM sucrose, 50 mM KCl, 2 mM KH_2_PO_4_, 20 mM Tris-HCl, 5 mM glutamate, 5 mM malate, pH 7.4, with additional 1.5 µM Calcium Green-5N and any compounds of interest, at a final concentration of 1.2 mg/mL. Plate was loaded into a PheraStarFX with injection ports pre-primed with CaCl_2_ at desired concentration. Assay was initiated by measuring fluorescence at 485_ex_/520_em_ (nm) and injections occurring at specified time-points for specific duration, depending on experimental set up. CRC/CRC_0_ ratios were calculated by determination of the injection cycle at which Calcium Green signal did not return below the spike point of injection, between treated and untreated mitochondria.

For bolus additions of CaCl_2_, mitochondria were aliquoted into 384-well plates at 4 mg/mL in 1.5 µM calcium green-5N, 130 mM sucrose, 50 mM KCl, 2.5 mM KH_2_PO_4_, 5 mM HEPES, 2.5 µM rotenone and 10 mM succinate (pH 7.4) and assay run as above, but with a single dose of calcium at the start of the assay. Quantitative analysis of bolus graphs was performed by using R (Version 4.0.0) by capturing the time taken for fluorescence signal to return to > 5 % of signal observed at minimum, with statistical comparison between treatment effects on wildtype and CypD KO mitochondria performed using analysis of variance. Replicates of the experiment were performed on different days, and day was included in the analysis as a block factor.

### Metabolic flux analysis

Oxygen consumption was measured using a Seahorse XF24 or XFe96 Extracellular Flux Analyzer (Agilent). For isolated mitochondrial studies, rat mitochondria were diluted in 220 mM mannitol, 70 mM sucrose, 5 mM KH_2_PO_4_, 5 mM MgCl_2_, 2 mM HEPES, 1 mM EGTA, 0.2 % fatty acid free BSA, pH 7.2 (mitochondrial assay solution, MAS) plus 10 mM succinate at 0.2 mg protein/mL. Mitochondrial suspension was added at 10 µg per well before centrifugation at 2000 x g for 20 min (4°C). 450 µL of MAS + 10 mM succinate, 2 µM rotenone, and any compounds of interest/vehicle were added to wells. Oxygen consumption was measured by the Seahorse XF Analyzer before and upon addition of ADP (5 mM), oligomycin (2 µM), FCCP (4 µM) and antimycin A (4 µM).

For cellular oxygen consumption assays, HEK293 cells were plated into Seahorse 96 well plates at 40,000 cells per well in DMEM media containing 10 % FBS. The next day, media was replaced with Seahorse DMEM media with 10 mM glucose, 1 mM pyruvate and 1 mM glutamine and mitochondrial stress test or glycolytic rate assay (Agilent) followed as per manufacturer’s instructions. Data was normalised by protein content measured using BCA assay after run completion.

Permeabilised cellular assays were performed using HEK293 cells treated with or without compound, permeabilised in MAS buffer containing 1 nM permeabilization agent (Agilent). Mitochondrial substrates glutamate and malate (10 mM each) were provided, and levels of oxygen consumption measured using a Seahorse XFe96 as above.

### Luminescence-based scintillation proximity assay

Human cyclophilin D sequence starting at amino acid 30, with an N-terminal 6His-TEV tag, was cloned into a pET24b plasmid between NdeI and HindIII sites. Resulting PET24b-His6TEV-CypD plasmid was transformed into BL21 (DE3) *Escherichia coli* and grown to 5 L in LB broth before pellet isolation, sonication-mediated lysis and isolation of protein by Ni-affinity and ion exchange chromatography. Assay and dilutions were all performed in 50 mM HEPES, 0.06 % Tween-20, 0.6 % BSA at pH 7.3. Purified cyclophilin D was pre-coupled to His-tagged copper yttrium silicate beads (Perkin Elmer) by incubation for 30 min on a rollerdeck at RT in the dark. CypD-bead complex (at a final concentration of 0.5 mg bead/150 nM cyclophilin D) and 10 nM [^3^H]-cyclosporin A (1 mCi/mL) were added to each well containing compound of interest. Mix was incubated on a plate-shaker at 750 rpm at RT for 30 min followed by centrifugation at 563 x g for 5 min at RT. Counts associated with [^3^H] were then measured using a TopCount (PerkinElmer) reading each well for [^3^H] over a 1-minute interval.

### Cell culture and transfection

HEK293 cells (ATCC) were maintained at subconfluency in DMEM (Gibco) containing 4.5 g/L glucose, 2 mM glutamine, 1 mM sodium pyruvate, 10 % FBS, with detachment performed using TrypLE express (Gibco). Cells were transfected with Lipofectamine 2000. Plasmid (CMV-NLRX1; Genscript) was combined with Lipofectamine for 30 min in OptiMEM (Gibco) before addition to a T175 flask containing HEK293 cells. Cells were incubated for a further 24 hours at 37 °C 5 % CO_2_ before media was replaced with fresh DMEM as above. Flasks were incubated for a further 48 h before mitochondrial isolation as above. Cell death in response to ionomycin addition was measured by activation of Caspase 3/7 according to manufacturer’s instructions (Promega).

THP-1 cells were obtained from ATCC and grown in RPMI + 10 % FBS. THP-1s were plated as required and incubated for 24 h with 20 ng/mL PMA. Media was removed and replaced for RPMI + 10 % FBS for a further 24 hours. Immune stimuli were then performed as suggested by Invivogen for 24 h.

Human primary macrophages were isolated from healthy human blood. PBMCs were isolated from human blood by density centrifugation with Ficoll. PBMC layer was resuspended in EasySep buffer and Monocyte Isolation kit (Stemcell Technologies) used to negatively select monocytes. Monocytes were plated into 96-well plates with 20 ng/mL M-CSF for 5 days, with media change at day 3. Macrophages were then exposed to immune stimuli as above.

### Imaging

mPTP imaging experiments were based on published methods ^25^. Cells were plated into single-chamber glass slides and left to adhere for 24 hours at 37 °C. Cells were washed in HBSS buffer containing HEPES (10 mM), L-glutamine (2 mM) and succinate (100 µM). Cells were then incubated in the buffer above containing 0.3 µM calcein AM, 1 µM MitoTracker CMX Red, 1 µM Hoechst and any additional compound for 10 minutes at 37 °C in the dark. Addition of 1 mM cobalt chloride was made, followed by further incubation for 5 minutes. Imaging buffer containing stains was removed and replaced with fresh buffer as above. Cells were imaged at the start of the assay using a Zeiss LSM880 AxioObserver with a Plan-Apochromat 63x oil objective with environmental control at 37 °C. Addition of 1 µM ionomycin was made and cells imaged after 12 minutes using the same settings as at the assay start.

For mitochondrial localisation, HEK293 cells were transfected with Lentiviral particles at an MOI of 5 for 72 hours before staining with 1 μM Mitotracker Red. Images were taken using a Zeiss LSM880 AxioObserver with a Plan-Apochromat 63x oil objective with environmental control at 37 °C before addition, and at 1h and 4 h after addition of DMSO (0.1 % v/v) or 10 μM GSK900.

Images were analyzed using Zen Lite and ImageJ software, including CoLoc2 for colocalisation quantification.

### Brain homogenate calcium retention assay

Mouse brains were harvested via decapitation and placed on ice. Brains were homogenised (1:2) in 130 mM sucrose, 50 mM KCl, 2.5 mM KH_2_PO_4_, 5 mM HEPES, 5 µM rotenone, 20 mM sodium succinate using a Dounce-homogeniser. Brain homogenates were spiked with 1 mM CaCl_2_ and incubated at room temperature for 12 min, and centrifuged at 15800 x *g*, 21 °C for 5 min. Calcium Green-5N was diluted 1:100 in the above buffer and 5 µL of this detection mixture was added to 95 µL of the sample supernatant and fluorescence measured as above for CRC assays. For experiments where mice were pre-dosed with vehicle/compound, brains were harvested 1 hour after dose. For experiments where brains were incubated with compound after harvest, compounds were preincubated for 15 min before addition of CaCl_2_.

### Chemoproteomics

Affinity enrichment of proteins bound to GSK864 was performed by covalently linking the carboxylic acid-bearing compound to NHS-activated sepharose beads (GE Healthcare). Unlinked beads were washed 3 times in 10 volumes of DMSO and resuspended in a 1:1 slurry. To achieve a coupling density of 2 µmol/mL sepharose, 20 µL of a 100 mM stock of GSK864 and 20 µL of triethylamine were added to 1 mL of settled beads and the mixture was incubated with rotation at RT for 16 h. To block non-reacted NHS groups, 50 µL of aminoethanol, 100 µL of diisopropylethylamine, and 100 µL of PyBrop (Millipore) was added and the mixture was incubated with rotation at RT for 16 h. Coupling efficiency was confirmed by HPLC of supernatants. Blocked control beads were also produced in the absence of GSK864. HEK293 lysate was generated and a BCA assay was used to normalise supernatants to 5 mg/mL in DP-buffer containing 0.4% IGEPAL and 1 mL of lysate was co-incubated with 50 µL of beads for 1 h at 4 °C. Beads were washed 10 times with 1 mL of DP buffer containing 0.4% IGEPAL and proteins were eluted in 50 µL elution buffer (100 mM Tris-HCl, 10% glycerol, 2% SDS, and 50 mM DTT). For competition assays, cell lysates were preincubated with 50 µM of respective compounds for 5 min at RT then co-incubated with 50 µL of beads for 1 h at 4 °C. Proteomics data generated in these experiments is captured in Supplementary Tables 1 and 2.

### Immunoblotting

Liver was collected from wildtype or *Nlrx1*^-/-^ mice and prepared as per the mitochondrial isolation above. Cell pellets harvested from HEK293 or liver mitochondria pellets were resuspended in RIPA buffer containing HALT protease (Thermo). Lysates were centrifuged at 13000 x g for 15 min and supernatants were assessed for protein concentration using a BCA assay. Protein was resuspended in LDS loading sample buffer containing reducing agent (Thermo), heated to 85 °C for 5 min, and loaded into a 4-12 % BisTris precast SDS-PAGE gel (Novex). The gel was run at 180 V in MES running buffer and transferred onto PVDF membrane using an iBlot2 (Thermo). Membranes were blocked in Odyssey block buffer (LI-COR) for 1 hour at RT and probed with rabbit anti-NLRX1 (Cell Signaling 1:1000), mouse anti-CypD (Abcam; 1:1000), mouse anti-VDAC (Abcam; 1:5000) or mouse anti-β actin (Sigma 1:10000) overnight at 4 °C. Membranes were washed 3 x 5 min in TBST before and after incubation with donkey anti-mouse or anti-rabbit secondary antibody (LI-COR) for 1 hour at RT, then imaged on an Odyssey CLX system.

### Protein concentration determination

Samples were assessed for protein concentration using a BCA assay according to manufacturer’s instructions (Pierce) by measuring absorbance of samples at 562 nm after 30 minutes incubation at 37 °C and comparing to known BSA standards.

### *In vivo* studies of GSK900

Mice were dosed *per os (p.o)* with GSK900 or 1 % methylcellulose at 5 mL/kg. Blood samples from tail vein or tissue samples were obtained at relevant time points. Samples were then analyzed using an API4000 (SciEx) Triple Quadrupole LC-MS/MS mass spectrometer (AB SciEx Instruments) in positive mode. To estimate the *in vivo* concentrations of free unbound compounds, protein binding was determined in blood and brain samples via equilibrium dialysis. Fresh pooled Wistar Han Rat blood was diluted with one volume of PBS (1:1, v/v) and brain was homogenised with two volumes of PBS (1:2, w/v). EDTA-K2 was used as anticoagulant. Compound was prepared at a target concentration of 5 µM in diluted blood and brain homogenate. The spiked blood and brain homogenates were placed on the donor side of the dialysis membrane (MWCO, 12-14K), and dialyzed against compound-free PBS by incubating the HTDialysis plate for 4 hours at 37 °C to achieve equilibrium. The concentrations of compounds on both sides of the membrane were analyzed using LC-MS/MS with subsequent values used to calculate the fraction unbound in blood and brain.

In vivo kainic acid induction of seizure modelling was performed by Transpharmation (UK). Mice submitted to the study were suitably blinded to genotype or treatment regime. Mice were treated *p.o* with vehicle (1% methylcellulose) or GSK900 (5, 30, 100 mg/kg, 10 mL/kg) 1 hour prior to induction of seizures with kainic acid (dosed s.c at 20 mg/kg, 5 mL/kg). Mice were monitored on the Racine scale for 180 minutes to record the time of first seizure, latency of seizure onset (Stage 3, 4, 5), number of animals manifesting each Racine stage stage 3 or above, total number and duration of each Racine stage over 3 and maximum seizure severity. Racine scale table is below. Mice were euthanised by cervical dislocation when they reached a humane endpoint (stage 5 continuously for 1 minute, stage 5 in brief but repeated bouts for 10 min or stage 6, or at the end of the 180 observation period.

| Racine Scale | Observed Behaviours |
| --- | --- |
| 1 | Facial clonus, staring, immobility |
| 2 | Rigidity, head nodding, blinking, tail extension |
| 3 | Forelimb clonus |
| 4 | Forelimb clonus with rearing |
| 5 | Rearing and falling, uncontrolled running and jumping, clonic seizures |
| 6 | Continuous or severe tonic-clonic seizures |

### CRISPR-Cas9 editing

CRISPR-Cas9 editing was performed using ribonucleoprotein delivery (Lonza). For *Ppif*, a single guide (IDT) was used and for *Nlrx1*, a triple guide approach (Synthego) was used. For *Nlrx1*, 25 µM of guide suspension was added to 10 µg of Cas9 protein in a total volume of 4 µL and incubated for 10 min at RT. 200,000 cells were resuspended in solution SF (Lonza) containing the RNP and electroporated on an Amaxa 4D using program CM130. For editing of *Ppif*, 0.1 crRNA, 0.1 nmol tracrRNA (IDT) and 3 µL duplex buffer (IDT) were assembled into a complex by heating to 95 °C for 5 min. Once cool, 10 µg of Cas9 protein was added and incubated for 15 min at 21 °C. Electroporation was carried out as above.

After electroporation, cells were rested for 48 hours at 37 °C, 5 % CO_2_. Editing success was measured by genotyping as described below. Successfully edited populations were diluted to single cell per well concentrations and grown into monoclonal colonies. Once confluent, cell colonies were assessed for genotype as described below and protein absence by immunoblotting as above.

### Genotyping

Cells were harvested by centrifugation at 300 x g for 5 min. Cell pellet was resuspended in 1 mM CaCl_2_, 3 mM MgCl_2_, 10 mM Tris, 1 mM EDTA, 1 % TX100, 0.2 mg/mL proteinase K, pH 7.4 and heated to 65 °C for 15 min then 95 °C for 10 min. PCR was run to amplify specified amplicon regions around the edit site. PCR mixture contained Phusion Mastermix (Thermo), 0.5 µM forward primer, 0.5 µM reverse primer, genomic material (50-250 ng) and water to 20 µL. PCR products were confirmed by agarose gel electrophoresis. PCR products were then purified by Qiagen PCR Purification kit (bulked edited populations) or PureLink 96-well PCR clean up kit (Thermo). Products were diluted to 2 ng/µL and sequenced with sequencing primers (GeneWiz). Sequencing results were analyzed using Synthego ICE tool with a wildtype control as baseline.

### ROS production

ROS production was measured in isolated HEK293 mitochondria aliquoted into 1536 well plates at 0.5 mg/mL. Mitochondria were incubated in 1 U/mL horseradish peroxidase, 50 µM Amplex Red and compounds of interest. Rotenone was used as a control at 300 nM to induce ROS production. ROS production, as measured by resorufin production, was measured by observing kinetic changes in fluorescence at 570_ex_/585_em_ (nm).

### Co-immunoprecipitation

Co-immunoprecipitation was performed by lysing HEK293 cells (WT or *NLRX1*^-/-^) that had been pre-incubated with DMSO or GSK900 for 4 hours, in RIPA buffer containing protease inhibitors for 15 min on ice. Lysate was clarified by centrifugation at 20000 x g for 20 min at 4 °C. Supernatant was then incubated with 1/50 dilution of anti-NLRX1 antibody overnight with gentle rotation. Protein A/G beads were used at 25 µL starting bead volume per sample in TBS + 0.1 % Tween-20. Isolated beads were resuspended in cell lysate containing antibody for 2 h at RT with gentle agitation. Beads were then isolated from the lysate using a magnet and uncaptured fraction stored for further analysis. Beads were then washed three times in TBS-Tween, before a final wash in water. Beads were then isolated on the magnet and resuspended in 50 µL 50 mM TEAB + 5 % (w/v) SDS, with 10 min sonication followed by 10 min incubation at 56 °C. Beads were then isolated on the magnet and remaining captured fraction stored at −80 °C for SDS-PAGE and proteomic analysis.

### Proteomics

#### Sample preparation

Cell pellets and isolated mitochondrial samples were defrosted at RT for 20 min and resuspended in 1x S-Trap lysis buffer (5% SDS, 50 mM TEAB) that was supplemented with Benzonase at 5000:1 dilution. The lysate was vortexed and mixed for 15 mins at 1500 RPM, RT. The lysate was centrifuged at 10000 x g for 10 minutes and a protein concentration quantified using a Pierce BCA Gold protein assay kit according to the manufacturer’s instructions.

For each sample, 10 µg protein was reduced with addition of TCEP added (10 mM final concentration) for 20 min at 60°C. Samples were subsequently carbamidomethylated for 30 min in the dark at RT by addition of iodoacetamide to a final concentration of 10 mM. Samples were acidified by addition of phosphoric acid to a final concentration of 1.2% before subsequent addition of 350 µL of S-Trap binding buffer (50 mM TEAB, 90% MeOH, pH 7.1). Samples were loaded onto a 96-well S-Trap™ plate (ProtiFi) and centrifuged for 2 min at 1500 xg. The columns were washed three times with 200 µL S-Trap binding buffer per well. Trypsin/LysC mix (Pierce) was prepared at a 1:25 sample to enzyme ratio in 50 mM TEAB and 125 µL added to each sample column. The plate was loosely covered and incubated for 2 h at 47°C. The peptides were eluted from the column by sequential addition of 80 µL 50 mM TEAB, 80 µL elution buffer 1 and 80 µL elution buffer 2 with 1500 xg centrifugations steps for 2 min. The peptides were then dried in a vacuum concentrator.

#### EvoSep tims-TOF Pro2

The Evosep One was coupled online to a hybrid (trapped ion mobility spectrometry) TIMS quadrupole TOF (time of flight) mass spectrometer (Bruker timsTOF Pro 2) via a captive spray nano-electrospray ion source. Samples were separated using a 60 SPD EvoSep method with an EV1109 performance column (8 cm x 150 µm, 1.5 µm). An ion mobility range was set from 1/K0 = 1.6 to 0.6 V s/cm^2^ with equal ion accumulation time and ramp times were applied in the dual TIMS analyser of 100 ms each. The ion mobility dimension was calibrated with the in-batch calibration function enabled. Mass spectra were recorded from 100-1700 *m/z* in positive dia-PASEF scan mode. When operating the mass spectrometer in diaPASEF mode, 8 diaPASEF scans per TIMS-MS scan were used, giving a cycle time of 0.95 seconds. For the diaPASEF windows, variable ion mobility windows (0.6-1.60 1/K0) and variable mass windows (300.2-1199.6 *m/z*) were generated using py_diAID. ^26^

### Data analysis

Raw mass spectrometry data files were analysed using DIA-NN (version 1.8.1).^27^ A library-free search was performed with a UniProt Human SwissProt only database (downloaded 2023). The following search parameters were used: peptide lengths of 7-30 amino acids with up to one missed cleavage, one fixed modification (carbamidomethylation; cysteine) and N-term M excision as a variable modification. Match between runs was enabled with a precursor FDR set at 1%.

Protein group files were analysed using Perseus 1.6.15.0 to perform filtering, transformation and statistical analyses. Gene ontologies were extracted using Enrichr and interaction analysis performed with STRING v.12. Data were visualised using GraphPad Prism v.8.0.

### Mitophagy assay

ARPE-19 (ATCC, CRL-2302) cells were maintained in 1:1 DMEM:F-12 media supplemented with 10% (v/v) FBS, 2 mM L-glutamine, 100 U/ml penicillin and 0.1 mg/ml streptomycin. The retroviral expression vector for *mito*-QC was previously described (Allen et al, 2013)^28^.

ARPE-19 cells stably expressing the *mito*-QC reporter were seeded in a 6 cm dish and treated with 10 μM of compound and 20 μM CCCP for 24 hours. After treatment, cells were washed with PBS, trypsinised for 5 min and centrifuged 3 min at 1,200 rpm. The pellet of cells was resuspended in 250 μL of PBS and 2 mL of 3.7 % (w/v) formaldehyde, 200 mM HEPES pH 7.0 were added. After 30 min at RT, 3 mL of PBS was added before centrifugation 5 min at 1,200 rpm. Finally, the pellet of cells was resuspended in 1% FCS in PBS and directly analysed by flow cytometry (as previously described^29^). Briefly, for each independent experiment, at least 5 × 10^4^ cells were acquired on LSRFortessa cell analyser. Based on FCS and SSC profiles, living cells were gated. To quantify the percentage of cells undergoing mitophagy, the ratio GFP/mCherry was analysed. The gate used for the control condition (DMSO) was applied to all the other conditions. The value used for this was based on quantitation of microscopy data from *mito*-QC cells that showed around 7 % *mito*-QC of cells had red-only puncta above the value of the mean.

### Structural modelling

Interactions between NLRX1 and small molecules were determined using AutoDock Vina following the methodology described by Jewell *et al,* and using previous modelling information^70–72^. For each compound, ten poses were generated and binding affinity scores were ranked from lowest to highest, with lower (more negative) scores corresponding to more favourable interactions. All figures were visualised and generated using PyMOL(TM) 3.1.6.1, Schrödinger, LLC.

### Quantification and statistical analysis

Unless otherwise stated, data are mean ± S.D. of at least three biological replicates. Two-condition comparisons used an unpaired two-tailed Student’s t-test. Seahorse parameters used two-way ANOVA with Bonferroni correction; where indicated, comparisons relative to vehicle used two-way ANOVA with Tukey correction. Significance: *P<0.05, **P<0.01, ***P<0.001, ****P<0.0001. Bolus CRC traces were quantified in R (v4.0.0) as the time for extramitochondrial calcium to return to >5% above minimum, compared by analysis of variance with experimental day as a block factor. Proteomic significance used unpaired t-test (p<0.05) with >0.58 log2 fold-change. NLRX1-GFP/MitoTracker colocalisation was quantified with Coloc2 as Pearson’s R. Per-experiment statistical details are given in the figure legends.

## Results

### Identification of potent inhibitors of the mitochondrial permeability transition pore

Among the recently characterised novel classes of mPTP inhibitors, some of the most thoroughly validated tools belong to the isoxazole and cinnamic anilide groups ^5,30–33^. Combining the structure-activity relationships established in these reports with findings from previous high throughput screening efforts ^21^, we produced a series of novel drug-like molecules to optimise potency, metabolic stability and blood-brain barrier permeability. Two lead compounds, the isoxazole GSK900 and the cinnamic anilide GSK864, were selected for further development from initial screens due to potency, favourable physicochemical properties and considerable chemical differences from the existing mPTP inhibitor, CsA (Figure 1A). Both GSK900 and GSK864 showed inhibition of calcium-induced mitochondrial swelling (Figure 1B) and enhanced capacity for calcium retention (Figure 1C) as compared to CsA in isolated mouse liver mitochondria. GSK900 was prioritised as lead and GSK864 as back-up, with an inactive control (GSK686, Supp Figure 1A) showing negligible inhibition in swelling and CRC assays (Figure 1D, E, Supp Figure 1B). Direct comparison of CRC ratios, calculated by the dose at mPTP opening in treatment vs control, indicate GSK900 and GSK864 were significantly more potent than CsA at all doses tested, with GSK864 being slightly more potent than GSK900 at higher doses (Figure 1F). Investigations of metabolic safety highlighted that acute exposure to CsA induced a diminished basal and State 3 respiratory rate of isolated mitochondria, likely explained by the promiscuous binding of CsA to non-cyclophilin proteins^34^, whereas GSK900 did not affect metabolic state in an acute setting (Supp Figure 1C). Importantly, GSK900 was not able to chelate calcium (Supp Figure 1D), indicating that its mechanism of inhibiting calcium-induced calcium-release is dependent on one or more biological targets and not direct calcium reduction, nor was GSK900 able to acutely induce ROS production in isolated mitochondria at concentrations used for mPTP inhibition (Supp Figure 1E, F). In addition, GSK900 was able to inhibit calcium-induced calcium-release via the mPTP in mitochondria isolated from human HEK293 cells (Figure 1G, 1H). Together, these data establish GSK900 as a potent, non-mitotoxic mPTP inhibitor in mouse and human mitochondria.

**Figure 1.**
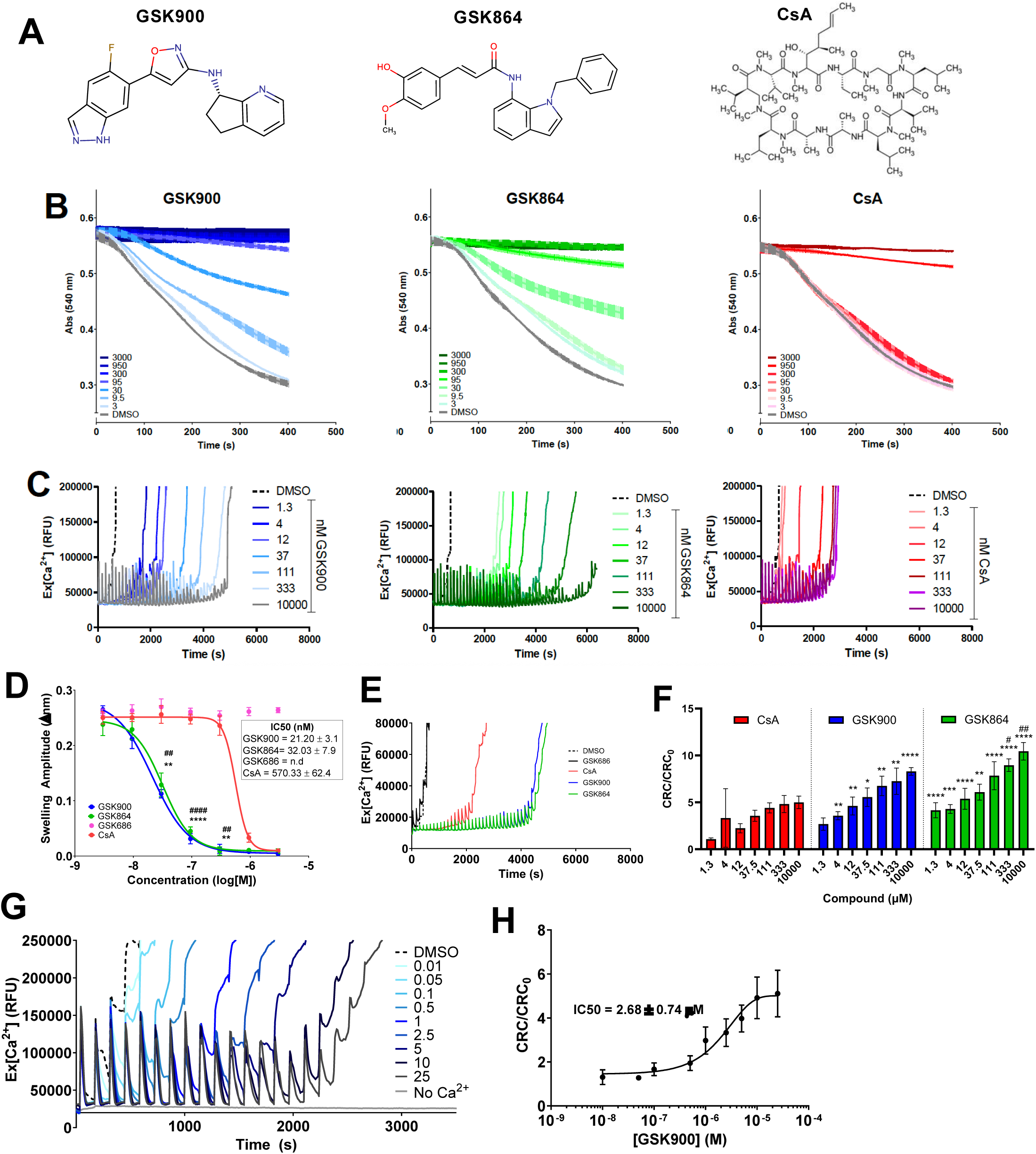
The compounds GSK864 and GSK900 are potent inhibitors of the mPTP in isolated mitochondria. **A**) Chemical structures of GSK900, GSK864 and CsA. **B**) Mouse liver mitochondrial swelling following addition of 800 nmol CaCl_2_ per mg protein in the presence of GSK900 (blue), GSK864 (green) or CsA (red) at indicated concentration. **C**) Mouse liver mitochondrial CRC upon incremental additions of 2 nmol CaCl_2_ per mg protein in the presence of GSK900 (blue), GSK864 (green) or CsA (red) at indicated concentration. **D**) IC50 of compounds in **(B)** and (**SF1B**) *;GSK900 vs CsA, #; GSK864 vs CsA **E)** Comparative CRC of compounds at 10 μM as in (**C). F**) Calculated CRC/CRC_0_ ratio across the dose range used in (**C)**. *; GSK900 or GSK864 vs CsA, #; GSK864 vs GSK900 **G**) HEK293 mitochondrial CRC upon incremental injections of 11.8 nmol CaCl_2_ per mg protein preincubated with GSK900. **H**) CRC/CRC_0_ ratios calculated from **(G)**. **All)** data is at least N = 3 biological replicates with data presented as mean ± S.D. **(B)**, **(C)**, **(E)** and **(G)** are representative traces from at least three biological replicates, as plotted in **(D)**, **(F)** and **(H)**. * = *P*<0.05, ** = *P*<0.01, ***= *P*<0.001, ****= *P*<0.0001 (# are used in **D** and **F** in the same way as *) as determined by a two-way ANOVA with Bonferroni multiple comparisons **(D, F)**.

### The biological activity of GSK900 is not mediated by CypD

It is well established that CsA inhibits mPTP activity by interacting with CypD, which positively modulates the affinity of the mPTP to Ca^2+^ ^35^. A previous study of isoxazole-based mPTP inhibitors reported the activity of the compounds was synergistic with that of CsA, indicating that the relevant molecular target might be something other than CypD^19^. Consistent with this, we observed that GSK900 exhibited an additive inhibitory effect when combined with CsA in mitochondria isolated from HEK293 cells (Figure 2A), as well as in mitochondria from mouse liver, kidney and heart (Figure 2B, Supp Figure 2A). This synergistic effect may well be modulated by enhanced concentration of two same-site compounds so to test whether GSK900 was able to interact with the CsA site of CypD, we performed a [^3^H]-CsA based scintillation proximity assay on purified recombinant CypD protein (Figure 2C). By increasing the concentration of unlabeled CsA in the medium, we were able to fully eliminate [^3^H]-CsA binding to CypD, verifying the canonical interaction, whereas GSK900 was not able to outcompete [^3^H]-CsA, even at 1000-fold molar excess. This confirms that GSK900 either binds to a region of CypD distinctive from CsA or, more likely, binds to a different target altogether.

**Figure 2.**
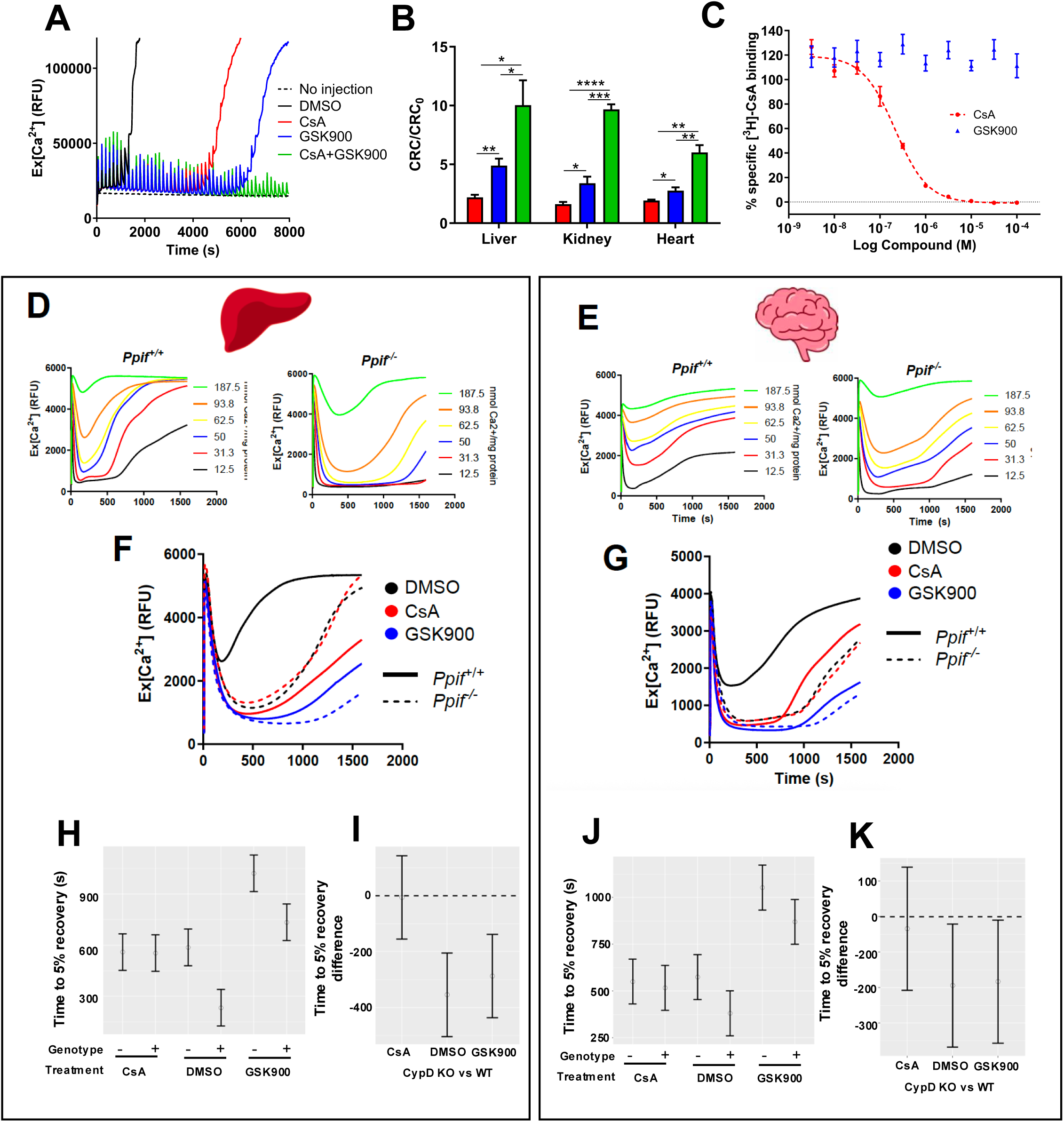
Influence of GSK900 on mitochondrial calcium permeability is independent of cyclophilin D. **A**) CRC of HEK293 mitochondria in the presence of 5 µM compound with 6 nmol CaCl_2_ per mg protein per injection. **B**) CRC/CRC_0_ ratios at 10 µM in mouse liver, kidney or heart mitochondria at 2 nmol CaCl_2_ per mg protein per injection. **C**) Binding of [^3^H]-CsA to purified CypD as measured by scintillation proximity assay in the presence of unlabelled CsA or GSK900. **D,E)** Measurement of calcium flux in wildtype or Ppif-/- mouse mitochondria from liver **(D)** or brain **(E)** upon bolus injection at indicated CaCl_2_ concentrations. **F,G)** Comparative analysis of fixed concentration of bolus CaCl_2_ to liver (**F**; 93.8 nmol per mg protein) or brain (**G;** 62.5 nmol per mg protein;) mitochondria from wildtype (solid lines) or Ppif-/-(dashed lines), pretreated with 0.1 % (v/v) DMSO (black), or 10 μM GSK900 or CsA 5 min prior to calcium addition. **H-K**) Quantitative analysis of traces in **F** and **G**, in liver **(H,I)** and brain **(J,K)** using mean duration from calcium addition (t=0) to mPTP opening, defined by an 5 % increase in fluorescent signal from minimum signal. Statistical comparison of the differences between wildtype and Ppif-/- in liver **(I)** and brain **(K)** mitochondria per treatment/genotype. **All)** Traces in **(A, D, E, F and G)** are averages of at least three biological replicates. Data is presented as mean ± S.D of at least three biological replicates. * = *P*<0.05, ** = *P*<0.01, ***= *P*<0.001, **** = *P*<0.0001 as determined by an unpaired two tailed T-test between conditions **(B)**.

To probe potential interactions between GSK900 and CypD, we utilised single-dose CRC assays in mitochondria isolated from wildtype or CypD KO (*Ppif*^-/-^) mice. Isolated liver (Figure 2D) and brain (Figure 2E) mitochondria were subjected to a variable bolus addition of calcium to induce mPTP opening. At all calcium doses, *Ppif^-/-^* liver and brain mitochondria showed delayed pore opening versus wildtype; CsA limited wildtype opening to *Ppif^-/-^* levels, whereas GSK900 inhibited more strongly than either CsA or *Ppif^-/-^* alone, and remained additive in *Ppif^-/-^* mitochondria. To enable quantitative interpretation of these bolus curves, the time taken for the extramitochondrial calcium signal to increase from its measured minimum by 5 % of (original signal – minimum), was calculated separately for each replicate in both liver (Figure 2H) and brain (Figure 2J). Using this analysis, CsA treatment, in either wildtype or *Ppif^-/-^*is no more effective in delaying mPTP opening than *Ppif^-/-^* alone, in both brain and liver, as expected. In contrast, GSK900 treatment delays mPTP opening longer than CsA treatment or *Ppif^-/-^*, but can also act in addition to *Ppif^-/-^* to further prolong mPTP opening. Contrasts to compare the two genotypes in each treatment statistically confirmed that CsA did not induce a difference in duration to mPTP opening between wildtype and *Ppif^-/-^*in either liver (Figure 2I) or brain (Figure 2K) yet GSK900 was able to significantly increase duration to opening in both. Together, these data confirm that the effect of GSK900 is additive to CsA in both human mitochondria and mouse mitochondria from several tissues, and is independent of CypD in its action.

### Identification of NLRX1 as a target for both isoxazole and cinnamic anilide mPTP inhibitors

Our finding that GSK900 acts via a CypD-independent mechanism is consistent with the behaviour of nearly all other non-peptide mPTP inhibitors reported to-date. To elucidate potential molecular targets for GSK900 we employed an affinity-based chemoproteomics approach that has been successfully applied to identify targets of other bioactive compounds identified in phenotypic drug screens (Figure 3A)^36–38^. In brief, an analogue of the active series is covalently coupled to beads, incubated with lysate in the presence of increasing concentrations of free compound, and captured proteins are quantified by LC-MS/MS; the free-compound IC50 for competed binding provides an apparent affinity for the compound-target interaction^39,40^.

**Figure 3.**
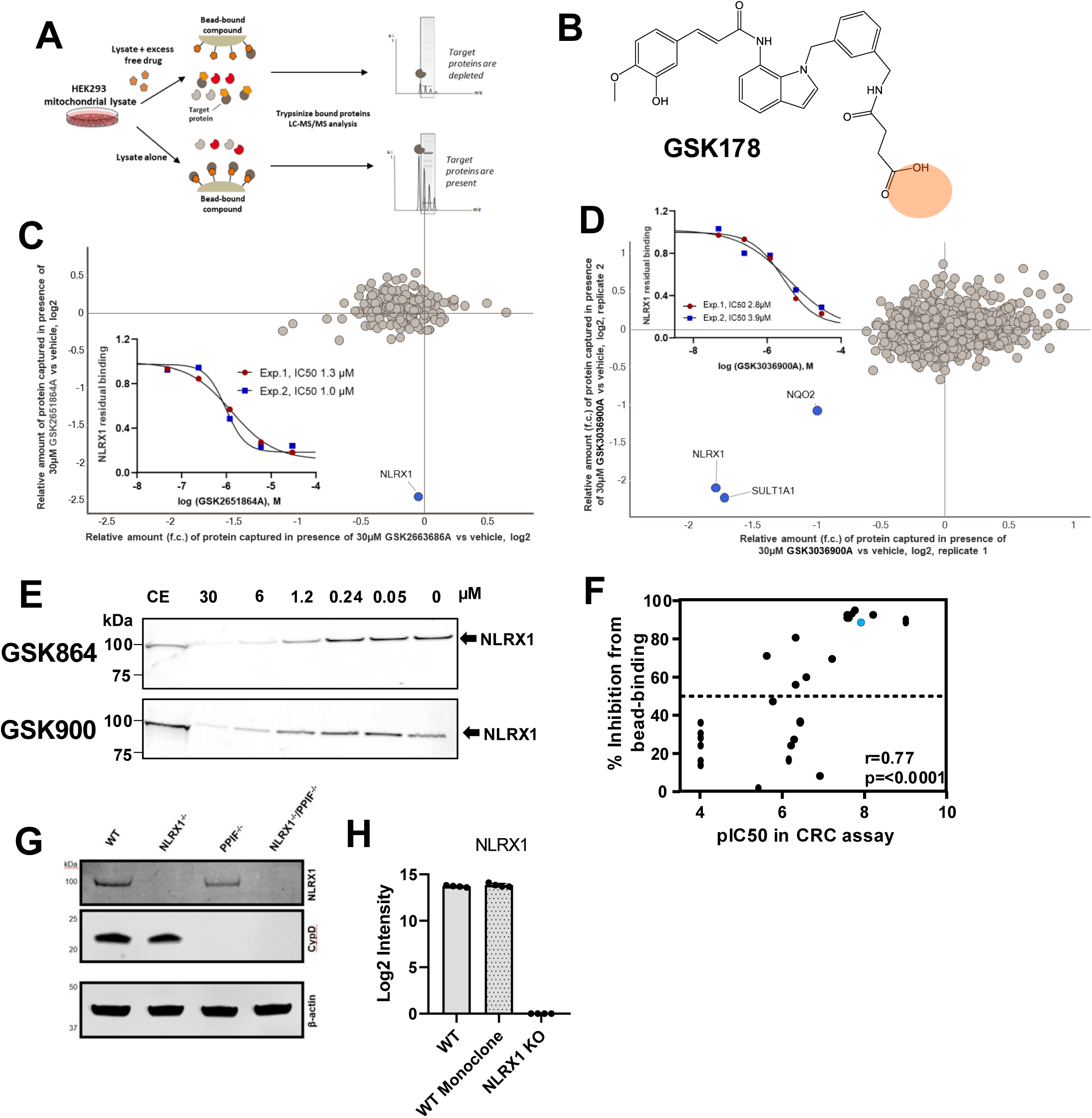
Chemoproteomic target deconvolution of GSK900 reveals NLRX1 as a biological target. **A)** Schematic of the chemoproteomics method where immobilised GSK178 is exposed to HEK293 mitochondrial lysate in the presence or absence of test compounds or vehicle (DMSO) for 1 h before capture, wash, elution and analysis by LC-MS/MS. **B)** Chemical structure of GSK178 **C)** Affinity purification +/- 30 µM GSK864 (active control; y axis) or GSK686 (inactive; x axis). The relative abundance of proteins captured by GSK178 in the presence of compound is expressed as a log2 fold change versus vehicle control. **Inset**; Determination of IC50 values for NLRX1 by competition binding experiments (as in **C**) in increasing concentrations of GSK864. **D)** Affinity purification +/- 30 µM GSK900. The relative abundance of proteins displaced by GSK900 versus vehicle was calculated and plotted for two independent experiments. **Inset**; Determination of IC50 values for NLRX1 by competition binding experiments (as in **D**) with increasing concentrations of GSK900. **E)** Western blot-based competition binding assay of NLRX1 to GSK178 +/- GSK900 or GSK864. **F)** Correlation of mitochondria CRC assay potency (pIC50) against % inhibition of NLRX1 binding at 10 µM as in (**C**) and (**D**) for 26 chemical analogues of GSK900 (blue). R; Pearson correlation coefficient, P; calculated probability. **G,H)** Western blot **(G)** or proteomic **(H)** confirmation of protein absence in *NLRX1^-/-^*, *PPIF^-/-^* or *NLRX1^-/-^/PPIF^-/-^* CRISPR-edited HEK293 cells.

As our lead compound, we attempted to functionalise the structure of GSK900 for use in these assays. However, we were unable to find a chemical variant that adequately maintained the lipophilicity and H-bond donors previously established as essential to mPTP-inhibitory activity. Our back-up molecule GSK864 possessed a similar IC50 to GSK900 in mPTP assays (Figure 1), but contains non-essential moieties that could be functionalised for covalent attachment to an agarose bead matrix. We therefore generated a functionalised version of GSK864 termed GSK178, by addition of a terminal carboxylic acid (Figure 3B). Due to the competitive nature of the assay, if GSK864 and GSK900 share common targets it would be possible to assess the depletion of such targets when GSK900 is present as “free” compound and monitor this depletion in a dose-dependent manner. Importantly, GSK178 exhibited similar biological potency to both GSK900 and GSK864 in swelling assays (Supp Figure2B, 2C). Through using direct comparison competition assays, it is possible to assess similar and different targets of the two biological classes of molecule, limiting the risk of false-negatives due to the lack of functionalised GSK900.

Having established that GSK900 was active in HEK293 experiments (Figure 2), we performed affinity pulldown experiments HEK293 mitochondrial lysates as depicted in Figure 3A. Incubation of lysates with the immobilised probe GSK178 was carried out in the presence of either DMSO as vehicle control and compared to first either incubation with an excess of GSK864 (Figure 3C, y axis), or an excess of the inactive analogue GSK686 (Figure 3C, x axis; all experiments were performed at least in duplicate and over a range of concentrations as described above). Following a wash step, the bead-bound proteins were trypsinised, subjected to LC-MS/MS analysis, and computationally aligned to the human proteome. We identified multiple proteins captured on GSK178-beads, but only NLRX1 was reduced on the beads in presence of excess active free GSK864, whereas we observed no reduction in incubations with inactive GSK686 (Figure 3C). GSK864 was shown to compete for binding of NLRX1 to GSK178-derivatised beads in a dose-dependent manner, with an IC50 of 1.0-1.3 μM (Figure 3C; inset).

Next, we assessed whether our original lead compound, GSK900, also targeted NLRX1 using a comparative competition assay. We performed affinity purification as above, with cross-competition by excess GSK900 (Figure 3D). Indeed, we observed that NLRX1 binding to GSK178-beads was reduced by addition of free GSK900 with an IC_50_ of 2.8-3.9 μM (Figure 3D), essentially confirming both GSK900 and GSK178 could bind NLRX1. We also identified two additional proteins interacting with GSK900, the cytosolic SULT1A1 with an IC_50_ of 5.5-8.7 μM and NQO2 with an IC_50_ of 13-15 μM. We reasoned that NLRX1 is the likely active target of both compounds as NQO2 is a cytosolic ubiquitously expressed redox-active protein which is a non-specific binder of several common drugs^41^, and SULT1A1 is cytosolic, a likely contaminant in this preparation, and our compounds were functional in isolated mitochondria.

To confirm the binding of GSK900 and GSK864 to NLRX1 in an orthogonal assay, we performed a Western Blot-based competition assay (Figure 3E). When cell lysates were incubated with GSK178-beads in the presence of increasing concentrations of GSK900 or GSK864, NLRX1 binding to GSK178-beads was inhibited in a dose-dependent manner. This assay was performed on 26 unique compounds from the original high-throughput screening (HTS) campaign, across both GSK900- and GSK864-related chemical series, selected from a range of pIC_50_ values (Supp Figure2D, E). Percentage reduction in NLRX1-binding to GSK178-beads in the presence of these compounds compared to vehicle control was seen to correlate with potency at the mPTP as determined in the HTS campaign, strongly indicating that NLRX1 binding is crucial for the biological activity of both chemical classes (Figure 3F, Supp Figure2D, 2E).

### NLRX1 is a positive regulator of mPTP function in human cells

To confirm the role of NLRX1 in the mPTP, we generated monoclonal NLRX1 KO (*NLRX1*^-/-^/*PPIF*^+/+^), CypD KO (*NLRX1*^+/+^/*PPIF*^-/-^) or double NLRX1/CypD KO (*NLRX1*^-/-^/*PPIF*^-/-^) HEK293 cells, and NLRX1 KO THP-1 cells by CRISPR-Cas9 (Supp Figure3A). Cell clones with suitable genetic edits as measured by sequencing were selected and protein absence was confirmed in the CRISPR-edited HEK293 and THP-1 cells by Western blot (Figure 3G, Supp Figure3B) and proteomics (Figure 3H).

Functionally, mitochondria isolated from wildtype (*PPIF^+/+^*;*NLRX1^+/+^*) and CypD KO (*PPIF^-/-^*; *NLRX1^+/+^*) cells underwent a typical mPTP response, albeit delayed in the *PPIF*^-/-^ setting, while GSK900 was able to dose-dependently delay mPTP opening in both genotype backgrounds (Figure 4A). Under these experimental parameters, genetic deletion of NLRX1 alone (*PPIF*^+/+^;*NLRX1*^-/-^) or in the CypD/NLRX1 double KO (*PPIF^-/-^NLRX1^-/-^*) from resulted in complete protection against mPTP opening (Figure 4A). To further investigate the efficacy of the compounds in intact cells, we used an established microscopy-based mPTP assay in HEK293 cells^25^. GSK900 was able to protect against ionomycin-induced mPTP opening in this assay, indicating that it is both cell penetrant and functional (Figure 4B-D). We also used this assay to investigate *NLRX1^-/-^* cells, which were also comparable to GSK900 or GSK864 treated wildtype cells, and CypD KO (*PPIF*^-/-^;*NLRX1^+/+^*) cells, whereas CsA-treatment was only partially effective (Figure 4C, 4D), and mitochondrial health remained intact over the course of the assay (Supp Figure3C, 3D). Overexpression of NLRX1 in HEK293 cells rendered mitochondria more sensitive to calcium induced mPTP opening, in a GSK900-sensitive manner (Figure 4E), suggestive that enhanced NLRX1 presence lowers the threshold for mPTP activation in a GSK900-sensitive manner.

**Figure 4.**
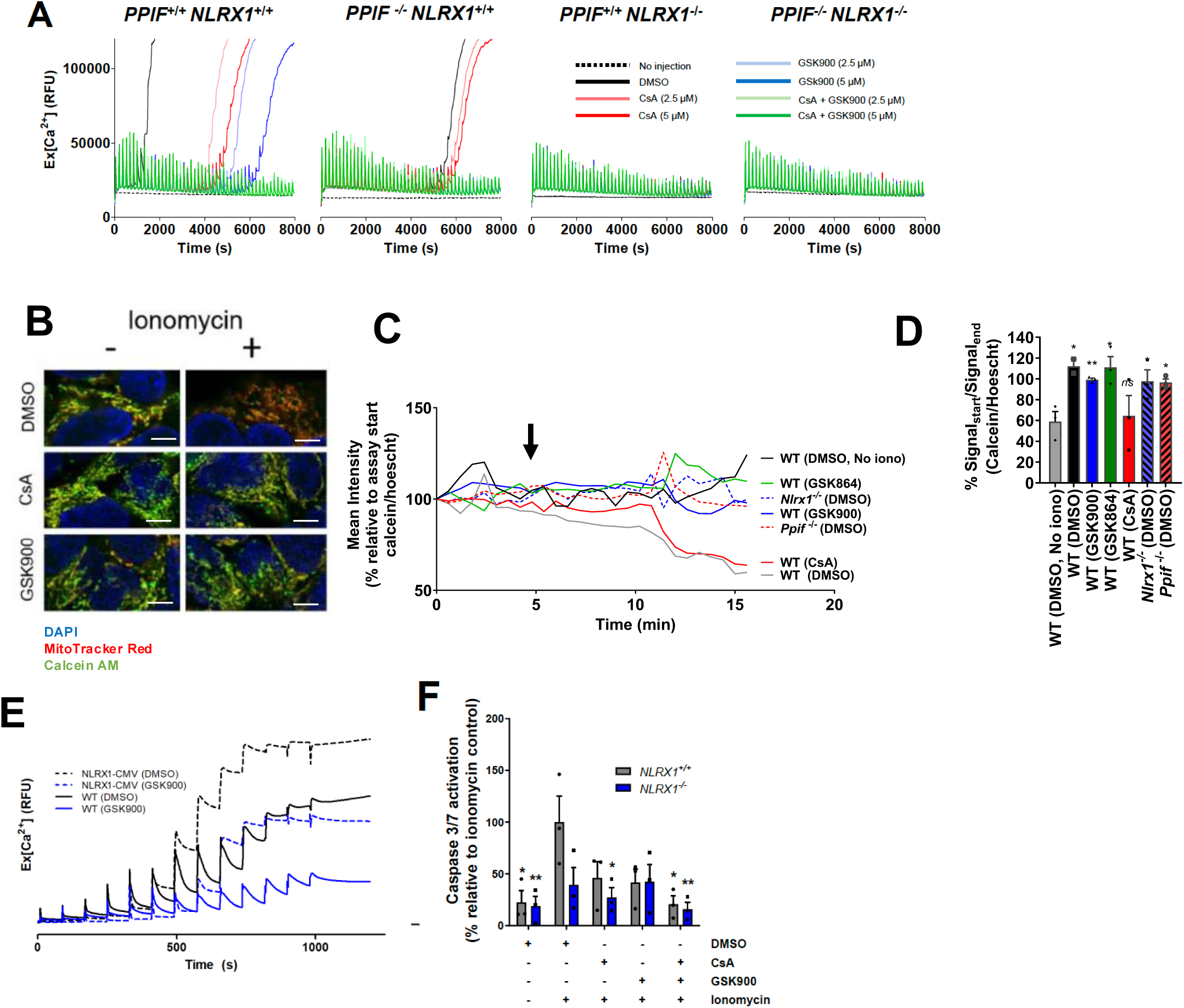
NLRX1 regulates the mPTP in human cells. **A)** CRC of HEK293 mitochondria isolated from WT or CRISPR-edited cells in the presence of 10 µM CaCl_2_ (6 nmol/mg protein). **B-D)** HEK293 cells pre-treated with calcein AM (green; 0.3 µM), MitoTracker Red (red; 1 µM), Hoescht 33342 (blue; 1 µM) and cobalt chloride (1 mM), and GSK900 or CsA (10 µM) before addition of ionomycin (1 µM) or vehicle (DMSO). Images in **(B)** represent at least three biological replicates, plotted kinetically in (**C)** and collectively in (**D)**, after 16 minutes of incubation with ionomycin. Scale bar in **(B)** represents 5 μM. **E)** CRC of NLRX1-CMV transfected (dashed) or wildtype (solid) HEK293 mitochondria with repeated additions of CaCl_2_ (7 nmol/mg protein), in the presence of DMSO or 5 µM GSK900. **F)** Ionomycin induced caspase 3/7 activation in wildtype or *NLRX1^-/-^* HEK293 cells, treated with 10 μM CsA or GSK900 for 30 min prior to 2.5 μM ionomycin addition. **All)** Data is presented as mean ± S.D of at least three biological replicates. * = *P*<0.05, ** = *P*<0.01, ***= *P*<0.001, **** = *P*<0.0001 as determined by a two-way ANOVA with Bonferroni multiple comparisons, except **(I)** where the statistics are relative to the vehicle condition (two-way ANOVA with Tukey’s multiple comparison test). Data in **(A), (C), (E)** are averages of three biological replicates with error not shown for clarity. Data in **(B)** and **(J)** are representative of at least 10 images, containing at least 5-10 cells per image. Each replicate in **(J)** is an average of all cells per image.

NLRX1 is the least-studied member of the NOD-like receptor protein family and the only member possessing an N-terminal mitochondrial targeting sequence. Physiologically, much has been learned from *Nlrx1*^-/-^ mice, which clarified a key role for NLRX1 in negatively regulating overactive innate immune responses through dampening of interferon and NF-κB signaling, as well as modulating cell death pathways and mitophagy^42,43, 44, 50^. GSK900 treatment or *NLRX1*^-/-^ was able to protect against ionomycin-induced cell death, as measured by Caspase 3/7 activation (Figure 4F). CsA treatment was partially additive to protection offered by *NLRX1^-/-^*whereas GSK900 was not in *NLRX1*^-/-^. GSK900 had only minor and inconsistent effects on the immune- and mitophagy-related functions previously ascribed to NLRX1, and did not phenocopy NLRX1 deletion in these assays (Supp Figure3J-M).

Altogether, these data highlight GSK900 has activity in human cellular scenarios with impact on mPTP, mitochondrial morphology and immune response, whilst not altering mitophagy and localisation in these tested conditions.

### Nlrx1 also regulates murine mPTP and GSK900 exhibits suitable *in vivo* pharmacokinetics and proof of concept efficacy

To explore this observation further we obtained *Nlrx1*^-/-^ mice which are viable and commonly used as a source of fibroblasts and bone-marrow derived macrophages^45^ and confirmed protein absence by Western Blot (Figure 5A). *Nlrx1^-/-^* mitochondria from heart, brain, liver and kidney exhibited a clear retention of mitochondrial calcium versus wildtype, however the kinetics of calcium uptake vs release differed in the brain and heart versus that in liver or kidney mitochondria (Figure 5B). Similarly, mitochondria isolated from *PPIF^+/+^NLRX1^-/-^* or *PPIF^-/-^NLRX1^-/-^* HEK293 cells also exhibited this trend in a repetitive high dose CRC assay, whereas WT or *PPIF^-/-^NLRX1^+/+^*mitochondria (Supp Figure3G), or WT treated with CsA (Supp Figure3H) or GSK900 (Supp Figure3I) behave as expected. This response could be explained by gradual accumulation of extramitochondrial calcium seen at these supraphysiological concentrations the uptake of Ca^2+^ via the mitochondrial calcium uniporter will likely become a limiting factor in the ability of mitochondria to completely sequester Ca^2+^ from the assay buffer.

**Figure 5.**
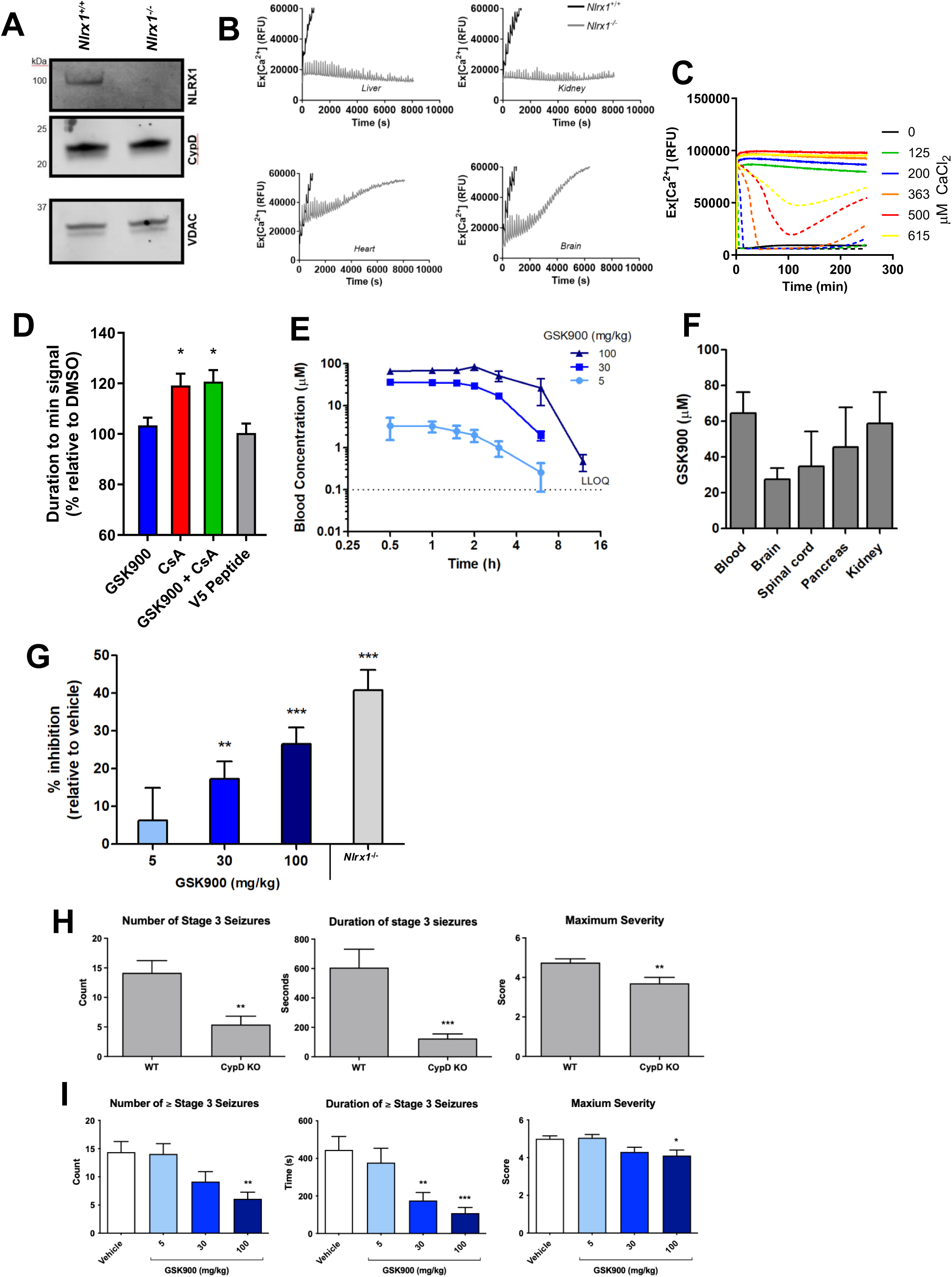
Nlrx1 regulates the mPTP in murine settings with brain penetrance and *in vivo* target engagement. **A)** Western blot confirmation of murine liver mitochondria from *Nlrx1*^-/-^ mice. **B)** CRC of isolated mouse liver, heart, kidney or brain mitochondria from wildtype or *Nlrx1^-/-^*mice, with repeat addition of CaCl_2_ at 2 nmol/mg protein. **C)** Calcium flux kinetics in liver mitochondria from wildtype (solid) or *Nlrx1^-/-^* (dashed) after single variable injection of indicated dose of CaCl_2_ at *t*=5s. **D)** Comparison of duration to mPTP opening as defined by inflection point of the curve set by the time where signal increased over 5% from minimum after single bolus CaCl_2_ addition (1.6 µmol/mg protein) in mitochondria isolated from *Nlrx1^-/-^*mouse liver in **(C)**. *; vs GSK900 **E)** Blood concentration of GSK900 over time following administration at 5, 30, or 100 mg/kg *p.o*. **F)** Tissue concentrations of GSK900 1 h following oral administration at 100 mg/kg. **G)** Calcium induced calcium release of brain homogenates from dosed, as indicated 1 h prior to culling, wildtype mice or *Nlrx1^-/-^* mice, and values normalised to calcium release measured in wildtype brain homogenates treated with vehicle (1 % methylcellulose). *; vs vehicle control. **H)** Kainic acid-induced seizure model in wildtype or Ppif-/- mice (n =>20). **I)** Kainic acid induced seizure model in wildtype mice pre-treated p.o with vehicle (1% methylcellulose) or GSK900 at noted dose for 1 hour prior to kainic acid injection (N>20 per group). **All)** data is presented as mean ± S.D of at least three biological replicates. * = *P*<0.05, ** = *P*<0.01, ***= *P*<0.001, as determined by an unpaired two tailed T-test between conditions. Data in **(B)** and **(C)** are averaged from three replicates with error removed for clarity.

To monitor supraphysiological mPTP activity in NLRX1 KO settings, we deployed the single bolus CRC assay (Figure 5C, 5D) against WT or *Nlrx1^-/-^*mitochondria. At these supraphysiological calcium concentrations, WT mitochondria were unable to sequester calcium and therefore do not exhibit a decrease in fluorescence associated with reducing extramitochondrial calcium, presumably due to immediate mPTP opening. In contrast, *Nlrx1*^-/-^ mitochondria were able to partially sequester the extramitochondrial calcium, exhibited by a decrease in fluorescence, yet with very high calcium addition did eventually release calcium in a dose-dependent manner (Figure 5C). Importantly, this calcium release observed in *Nlrx1^-/-^* mitochondria was delayed by CsA but not GSK900 or V5-inhibiting peptide, a Bax pore blocker, suggestive of CypD-sensitive mPTP activity (Figure 5D).

To preliminarily evaluate the *in vivo* pharmacology and biological effects of GSK900 we first characterised the pharmacokinetic (PK) profile in mice. Blood exposures (Figure 5E) after oral dosing at 5, 30, or 100 mg/kg and tissue exposures (Figure 5F) at 1h after a 100 mg/kg oral dose highlight stable and widespread dosing, including brain penetrance. The unbound fractions of GSK900 in blood and brain, estimated via *in vitro* equilibrium dialysis, were 0.72 ± 0.03 % and 1.31 ± 0.03 % respectively and therefore the free concentrations in brain are expected to exceed the IC50 concentration for GSK900 as measured in swelling assay at both 30 and 100 mg/kg dose levels. The oral bioavailability of GSK900 was 66 % and 61 % at 30 mg/kg and 100 mg/kg *p.o* doses, respectively, and when dosed intravenously at 1 mg/kg, exhibited a half-life of 0.64 h with a blood clearance rate of 18.67 mL/min/kg. To confirm observed brain penetrance, an *ex vivo* brain homogenate Ca^2+^ retention assay was utilised (Figure 5G). GSK900 dose-dependently reduced mPTP-mediated mitochondrial Ca^2+^ release following CaCl_2_ addition to brain homogenates harvested from mice 1 hour post dose, with 100 mg/kg dosing approaching inhibition levels seen in *Nlrx1^-/-^*mice.

As a preliminary *in vivo* efficacy study, we deployed the kainic acid seizure model as a proof of concept mPTP-driven neurological injury model. To provide confidence in the model, mice lacking CypD were first used to assess the potential window of mPTP-driven opportunity, with Ppif-/- mice exhibiting significantly reduced number, duration and severity of seizures (Figure 5H). Next, mice were dosed with 5, 30 or 100 mg/kg GSK900 1 hour prior to induction of seizures. GSK900 exhibited a dose-dependent reduction in number, duration and severity of seizures (Figure 5I).

These data provide preliminary pharmacological proof-of-concept that GSK900 reaches the brain and is active in an mPTP-sensitive neurological injury model

### NLRX1 associates with mPTP-linked mitochondrial proteins in a GSK900-sensitive manner

To investigate the mechanistic relationship between NLRX1 and the mPTP, we deployed investigative proteomics to assess global proteome and protein-protein interactions upon NLRX1 deletion and/or treatment with GSK900. Firstly, proteomes were obtained from whole cell and mitochondria, from WT and *NLRX1*^-/-^ HEK293 cells, using an established proteomic pipeline (Figure 6A), with suitable separation between GO and KEGG components between whole cell and mitochondrial proteomes (Supp Figure4A). Comparing WT and *NLRX1^-/-^* mitochondria (Figure 6B) highlighted 80 proteins downregulated in the *NLRX1^-/-^*setting. Network analysis of these downregulated proteins revealed modulation of pathways associated with mitochondrial respiration and protein insertion (Figure 6C). Next, we assessed the proteomic shift in whole cells upon treatment with GSK900 (Figure 6D). Whilst GSK900 treatment in WT cells caused a marked downregulation of proteins associated with mitochondrial function (Figure 6Di), addition of GSK900 to *NLRX1*^-/-^ HEK293 cells did not exhibit the same magnitude of downregulated proteins (Figure 6Dii). Interestingly, a correlation comparison between proteomes of *NLRX1^-/-^* cells and WT cells treated with GSK900, as compared to WT cells treated with vehicle, elucidated a shared downregulation of mitochondrial proteins, namely ATP synthesis and protein import (Figure 6E, Supp Figure4B).

**Figure 6.**
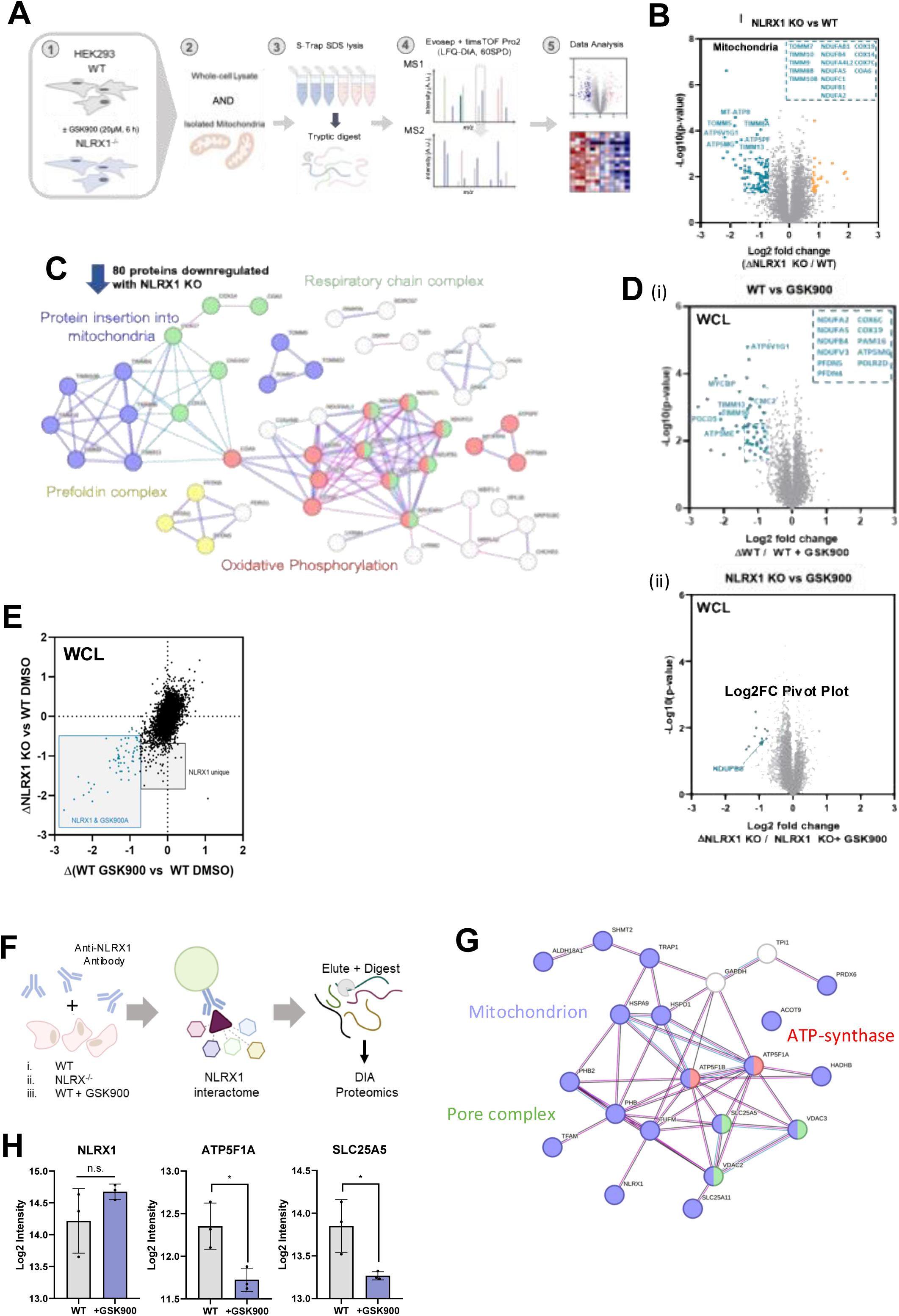
Investigative proteomics reveals NLRX1 modulation mediates a loss of mitochondrial proteins. **A)** Workflow for LFQ-DIA proteomics experiments 1) generation of treated samples, 2) isolation of mitochondria from WT & NLRX1^-/-^ treatments 3) SDS-Lysis and S-Trap digestion of samples 4) LFQ-DIA acquisition on an Evosep-timsTOF Pro 2 system 5) subsequent data analysis. **B)** Proteome of mitochondria isolated from wildtype and *NLRX1*^-/-^ HEK293 cells. **C)** interaction network of downregulated proteins represented as a STRING-DB network with relevant GOs highlighted. **D)** Volcano plots of wildtype **(i)** or *NLRX1*^-/-^ **(ii)** HEK293 cells treated with 20 μM GSK900 for 6 h. **E)** Correlation plot of Log2 intensities from wildtype cells treated with vehicle versus GSK900 (20 μM 6h) against wildtype and *NLRX1*^-/-^ cells (DMSO 6h) from **(D)**. Statistically significant proteins determined with an unpaired Students T-test (p-value <0.05) exhibiting >0.58 log2 fold change (lfc). **Co-immunoprecipitation of NLRX1 pulls out mPTP-interacting partners with GSK900 sensitivity (panels F-H)** **F)** Workflow for NLRX1 mediated co-immunoprecipitation and subsequent LFQ-DIA proteomic acquisition. **G)** STRING-DB representation of detected proteins captured by NLRX1-mediated co-immunoprecipitation. Proteins were excluded if 1) non-mitochondrial according to MitoCarta3.0 and 2) presence in a co-immunopreciptation performed using the same anti-NLRX1 antibody in lysates from *NLRX1^-/-^* lysates. **H)** Log2 intensities of select proteins from co-IP with treatment of GSK900. Statistically significant proteins determined with an unpaired Students T-test (p-value <0.05) exhibiting >0.58 log2 fold change (lfc) difference in abundance in pulled down samples between wildtype cells treated with vehicle or GSK900 (30 μM, 2 h). *=*P*<0.05.

As treatment with GSK900 phenocopied NLRX1 deletion, as measured by proteomics with a depletion of mitochondrial proteins, we next asked what interaction partners NLRX1 has, and whether this could explain the decrease in mitochondrial protein abundance in these scenarios. To do this, a coimmunoprecipitation of NLRX1 followed by proteomic assessment of the interacting proteins, or interactome (Figure 6F, Supp Figure5A), was performed in WT cells treated with DMSO or GSK900, and also *NLRX1^-/-^* cells as a negative filter for indirect interactions within the immunoprecipitation. Notably, alongside the known NLRX1 interactor TUFM, an enrichment of mitochondrial proteins was detected (Supp Figure5B). Of the detected mitochondrial proteins (Figure 6G), a number were associated with mPTP complex and ATP synthase. Upon treatment with GSK900, several proteins were depleted from the interactome but the only significantly reduced proteins were ATP5F1A, a subunit of the ATP synthase, and SLC25A5, a mitochondrial ATP/ADP translocase; both postulated mPTP components.

### NLRX1 modulation alters ATP synthase-associated proteome and mitochondrial respiration

Next we investigated the impact of GSK900 or *NLRX1^-/-^*on ATP synthase as one of the identified networks modulated by NLRX1 modulation in Figure 6, and as ATP5F1A was identified as an interaction partner modulated by GSK900 in Figure 6H. Notably, in the proteomics performed in Figure 6, a number of ATP synthase subunits or associated proteins were noted to be downregulated in *NLRX1^-/-^* cells and/or GSK900 treatment. Of those ATP synthase related proteins detected in proteomics, 6/13 were noted to be decreased with GSK900 treatment, and *NLRX1^-/-^*(Figure 7A, B). Several ATP synthase subunits (ATP5F1A, ATP5F1B, ATP5F1E and ATP5F1C) were noted to be decreased in *NLRX1^-/-^* cells, but not with GSK900 treatment, and ATP5F1D, ATP5MF and MT-ATP6 were not modulated by GSK900 treatment or NLRX1 deletion.

**Figure 7.**
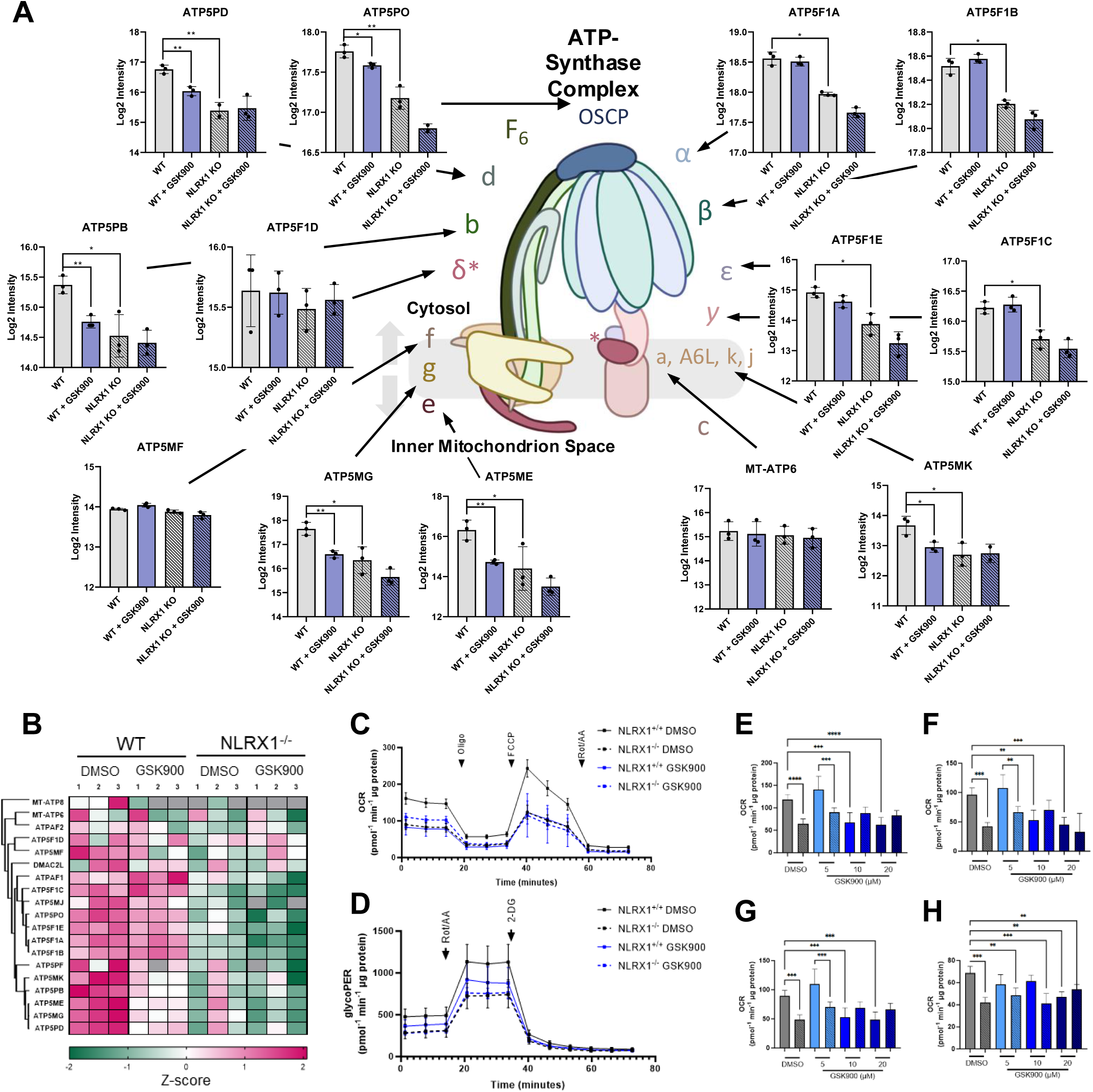
Chronic NLRX1 modulation mediates a loss of ATP synthase subunits and metabolic function. **A)** Abundance of proteomically-detected ATP synthase subunits in wildtype (solid) or *NLRX1^-/-^* (dashed) HEK293 cells treated with either DMSO (grey) or 20 μM GSK900 (blue) treatment for 6 h. **B)** Heatmap visualisation of ATP synthase subunits as in **(A)**. **C-G)** Seahorse analysis of wildtype (solid) or *NLRX1^-/-^*(dashed) HEK293 cells, treated with vehicle (black) or GSK900 (blue; 20 µM in **C, D**; as indicated in **E-H**) to indicated dose for 6h before in **C,D**) assessment by Mitochondrial Stress Test **(C)** or Glycolytic Rate Test **(D)** with selected parameters highlighted in **E-H; E)** Basal Respiration, **F)** Spare Respiratory Capacity **G)** ATP Production. **H)** Wildtype (solid) or *NLRX1^-/-^* (dashed) HEK293 cells treated with vehicle (grey) or GSK900 (blue) as indicated doses for 6 hours before permeabilisation and basal OCR assessment by Seahorse. **All)** Data is presented as mean ± S.D of at least three biological replicates.

We had previously assessed the metabolic impact of GSK900 on isolated mitochondria in an acute setting (Supp Figure1C), however given the above data in the context of longer-term exposure to GSK900 or in NLRX1 deletion settings, we revisited the functional metabolic flux analysis in a similar chronic setting (Figure 7, Supp Figure 6). Seahorse assays testing mitochondrial and glycolytic function were run on whole (Figure 7C-G) and permeabilised WT and *NLRX1^-/-^*HEK293 cells (Figure 7H). Cells treated with GSK900 or lacking NLRX1 exhibited diminished function, across several parameters in both mitochondrial function (Figure 7C) and glycolysis (Figure 7D), including basal respiration (Figure 7E), spare respiratory capacity (Figure 7F) and ATP production (Figure 7G), whilst treatment of *NLRX1^-/-^* cells with GSK900 had no additional impact on metabolic flux. In a permeabilised cell system, using succinate as a direct substrate to offset any potential metabolite transport conditions, NLRX1 deletion or GSK900 treatment induced the same blunted metabolic response as seen in whole cells (Figure 7H).

## Discussion

The physiological significance of the mPTP has long been appreciated, but an incomplete molecular identity has limited therapeutic targeting of the mPTP. Here we have incorporated learnings from published studies of non-peptide mPTP inhibitors to develop a brain-penetrant isoxazole mPTP inhibitor, GSK900, as well as a cinnamide-based GSK864. While an isoxazole compound was previously shown to improve motor function in a zebrafish model of muscular dystrophy^19^, ours is the first example of this chemotype with confirmed CNS target engagement and efficacy after oral dosing. Importantly, a biological target of GSK900 was elucidated as NLRX1, with impact in both murine and human settings associated with NLRX1 and mPTP function. Using proteomics, we were able to suggest a putative mechanism by which NLRX1 interacts in a GSK900-sensitive manner with the postulated mPTP proteins ATP5F1A and SLC25A5, and that in the presence of GSK900 or in settings of NLRX1 depletion, mitochondrial function is impacted due to depletion of key mitochondrial proteins such as ATP synthase subunits, protein import machinery and oxidative phosphorylation machinery. In addition, docking into a published hexameric homology model of NLRX1 (Supp Figure 7) offers hypothesis-generating clues to candidate binding sites and key residues, and is consistent with the observed structure-activity relationship: the hydrogen-bond-donating hydroxyl of GSK864, replaced by a methoxy in the inactive GSK686, may underlie the difference in activity between these analogues.

During the drafting of this manuscript, evidence implicating NLRX1 in the mPTP has emerged ^68, 69^. Xiao *et al.* has shown that NLRX1 is decreased in ischemia-reperfusion injury and that NLRX1 deletion abolishes calcium-induced mPTP opening in the heart, contributing to cardioprotection through control of metabolic pathways including mTOR, AMPK, and RISK signaling. Independently, an unbiased phenotypic CRISPR screen across more than 19,000 genes identified NLRX1 as the only mitochondrial hit whose deletion increased calcium retention capacity and abolished calcium release, meeting the authors’ definition of an essential activator of the permeability transition, while classical candidate pore proteins were dispensable. Our study reaches the same conclusion by a third, orthogonal route: chemical target-deconvolution from two structurally distinct inhibitor series. That cardiac genetics, unbiased human genetic screening and chemical biology independently converge on NLRX1 provides strong, mutually reinforcing evidence for its role as an essential regulator of mPTP opening. Uniquely, our work also provides brain-penetrant chemical matter with which to interrogate and pharmacologically modulate this biology. Collectively, these converging lines of evidence establish NLRX1 as an essential regulator of mPTP opening and open a route to indications where the mPTP is implicated, with our compounds providing tools to explore this biology.

The use of efficacious chemical matter coupled with chemoproteomics to identify a novel regulator of the mPTP is without precedent in the literature. Chemoproteomic identification of the GSK900-NLRX1 interaction (Figure 3) is not alone sufficient to establish NLRX1 as an mPTP regulator; however, our genetic validation in both CRISPR-edited cells (Figure 4) and *Nlrx1^-/-^* mouse tissue (Figure 5), and follow up proteomic-led investigations (Figure 6), establish that NLRX1 is an essential regulator of the permeability transition process; we interpret this as a functional requirement for normal calcium-induced pore opening rather than evidence that NLRX1 is a structural component of the pore, as well as a regulator of general mitochondrial homeostasis. Although functional genomic techniques will likely identify additional regulators in time, this study has provided hit identification in intact wildtype organelles, rather than selectively reconstituted systems^8^, as well as rapid translation using the same chemical matter to proof of concept use in a murine system. Obvious next steps would be to test GSK900 in the numerous disease models in which *Nlrx1*^-/-^ mice have shown protection, albeit with consideration to the impact on mitochondrial function observed here^9,42,48^. Additionally, the ability of GSK900 to compete with GSK864 for affinity matrix binding of NLRX1 suggests that the core functional group which distinguishes cinnamide from isoxazole compounds is dispensable for the interaction (Figure 3). Thus, NLRX1 is a CypD-independent (Figure 2) regulator of the mPTP and a shared biological target of both compound classes.

A central question raised by our study is how NLRX1 exerts its influence over the mPTP. In addition to suppression of anti-viral signalling via interactions with TUFM, RIG-I and MAVS on the OMM^45, 51^, NLRX1 can influence complex III-mediated ROS generation via UQCRC2 on the IMM^46,47^. NLRX1 trafficking and its N-terminal domain remain controversial, though we saw no effect of GSK900 on NLRX1 localisation, as well its cytosolic partners (TRAF6, LC3, caspase 8) are unlikely to act in purified mitochondria^49,52^.

Importantly, NLRX1 deletion did not alter calcium uptake, suggesting the NLRX1 acts only on the mPTP-mediated calcium release process, whilst overexpression studies show increased concentrations of NLRX1 are sufficient to decrease the threshold for mPTP activation. Our proteomic studies highlighted several previously unknown interaction partners for NLRX1, including ATP5F1A and SLC25A5 which exhibited significant GSK900-sensitivity in their interaction with NLRX1. Both proteins have been suggested as putative components of the mPTP, with some tissue dependency^53,54^. NLRX1 may exert its influence over the mPTP, and perhaps wider mitochondrial biology, through these molecular interactions.

Although NLRX1 deletion and GSK900 impacts mitochondrial protein abundance and function in chronic settings, GSK900 is still able to inhibit the mPTP in an acute setting, with almost immediate effect and is therefore unlikely to require a chronic depletion of mitochondrial proteins. For example, in mitochondrial CRC assays GSK900 was only present for a maximum of 5 minutes prior to calcium addition, suggesting the biological efficacy of GSK900 on the mPTP is not dependent on modulation of the proteome. We therefore propose that NLRX1 acts on two distinct timescales: an acute, proteome-independent role in setting the calcium threshold for pore opening (evident within minutes in CRC assays), and a slower, homeostatic role in maintaining mitochondrial protein content (evident over hours). On this model, the global proteomic changes are a secondary consequence of disrupting NLRX1 and other interaction partners, i.e ATP synthase. The phenocopying nature of the pharmacological and genetic modulation of NLRX1 highlights that removal of NLRX1, in either fashion, ultimately culminates in a diminished mitochondrial phenotype, elucidating a critical role for NLRX1 in mitochondrial homeostasis in longer term settings. This may have relevance to linking metabolism to immune response, and NLRX1 has been recently linked to immunometabolism of CD4^+^ T cells, which may itself be determined by the function of NLRX1 disclosed here^55,56^, through interactions with other protein families known for balancing mitochondrial homeostasis^55^, and mitophagy^43^. The chronic proteomic depletion may partly reflect in vitro conditions, so mitochondrial function should be monitored in longer-term in vivo dosing. Reassuringly, *Nlrx1^-/-^*mice are viable with no major safety issues in breeding or in our (short-term) dosing, suggesting NLRX1-targeting molecules are unlikely to be overtly detrimental *in vivo*.

Whether NLRX1 is itself an integral part of the pore remains open. Distantly related plant NLRs can directly form ion-permeable pores, and although NLRX1 has never been shown to do so, a direct structural contribution cannot be formally excluded here^57^. We treat this as an open possibility rather than a claim: our functional data establish NLRX1 as required for pore opening but do not demonstrate it is a physical constituent of the pore. *Nlrx1*^-/-^ mitochondria were still able to release calcium back into the extramitochondrial environment, which was delayed by additional CsA which does not affect calcium uptake. On its face this indicates that a CsA-sensitive pore can still form in the absence of NLRX1, albeit severely delayed and only under extreme, supraphysiological calcium loads. We note, however, that this observation sits in apparent tension with the recent finding that NLRX1 deletion abolishes calcium release in a CsA-insensitive manner in a cell-based permeability assay. We do not attempt to resolve this discrepancy here, and flag it as an open question: the difference may reflect the distinct assay regimes (isolated-organelle bolus calcium-retention measurements versus a cellular inner-membrane permeability readout), residual calcium handling at supraphysiological load, or a genuine difference in the residual pore, and distinguishing these will require further dedicated study. Either way, our data are consistent with NLRX1 being required for normal, physiological-threshold pore opening rather than for pore formation per se.

Considering the role of mitochondrial homeostasis in regulated cell death, a positive regulator of the mPTP might be expected to play a critical function in cell death processes. NLRX1 has been linked to both pro- and anti-apoptotic functions^58–62^, and the mPTP is permeable to biomolecules up to 1.5 kDa, including mtDNA fragments and formylated peptides; NLRX1 may thus regulate innate immunity in part by controlling mPTP-mediated release of such mitochondrial DAMPs, which activate sensors such as cGAS under oxidative or calcium stress^63–67^.

Altogether, using chemoproteomics, genetic validation and investigative proteomics, this study conclusively links NLRX1 to mPTP biology and mitochondrial homeostasis. These findings shed light on the atypical mitochondrial localisation of NLRX1 and suggest that NLRX1 may exert its influence on cellular biology, and in particular immunity, through direct control of the mPTP and mitochondria during times of biological stress, such as infection. This work connects innate immunity and mitochondrial biology through NLRX1, the least-studied NOD-like receptor.

## Author contributions

L.M.B, R.P-H, R.J.P, K.S, J.C.B, K.T, M.T, L.P.W, O.W, S.R, S.G-D, Y-M.T, W.X, C-C.A, E.R, G.B, A.J.B, D.G, performed experimental work and P.O, E.R, R.G.P, Y.L, K.C.L performed chemical synthesis, with project supervision from L.M.B, E.B, A.B, G.D, M.A. D.W.F co-wrote the manuscript and advised on the project. I.G.G and L.P.W led and performed the mitophagy assay, respectively. K.H managed *in vivo* work. N.G assisted with quantification and statistical interpretation. A.G, A.R and K.S provide structural modelling expertise. L.M.B, D.W.F and R.P-H wrote the manuscript with input from all authors.

## Acknowledgements

We would like to acknowledge all those scientists and staff that worked on this project over the years. In particular, the support of the Immunology Network and its leaders over the years for sponsoring and funding several studentships, postdoctoral fellowships and secondments that all contributed to this project. A special mention to Fiona Reed (GSK) for her operational support on this program. In addition, to several academic collaborators that helped shape and guide this project. Support from the DSTT-GSK collaboration (I.G.G, L.P.W) aided the mitophagy experiments.

## Data and Material Availability Statement

Proteomics datasets have been uploaded to PRIDE with identifier; PXD075000. Please contact corresponding author for requests.

## Supplementary Figures

**Supplementary Figure 1:**
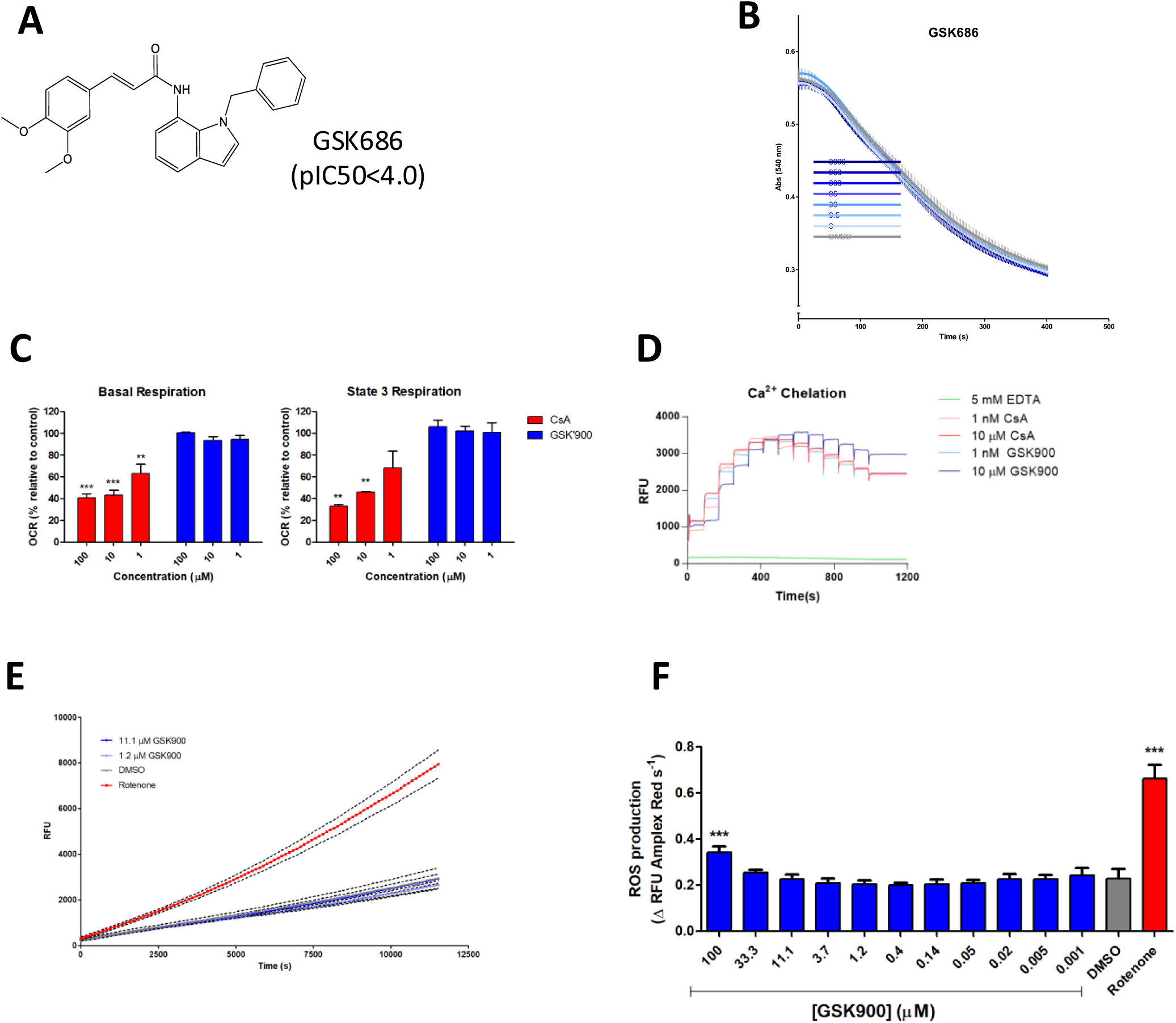
**A)** Structure of GSK686, inactive control. **B)** Mitochondrial swelling assay of GSK686, as in Figure 1B. **C)** Basal and State 3 Respiration of isolated rat liver mitochondria exposed to indicated doses of CsA or GSK900 for 15 min prior to start of assay. **D)** Calcium injections of 2 nmol in the presence of CsA (red) or GSK (blue) and EDTA (green) to assess calcium chelating potential. **E, F)** ROS production of rat liver mitochondria as measured by amplex red assay, plotted kinetically in **(E)** and as Δ RFU over the course of the assay **(F)**. **=P<0.01, ***=P<0.001 as determined by an unpaired two tailed T-test between conditions. Data is presented as mean ± S.D of at least three biological replicates. Data in **(B)** and **(E)** is average of three replicates with error omitted for clarity.

**Supplementary Figure 2:**
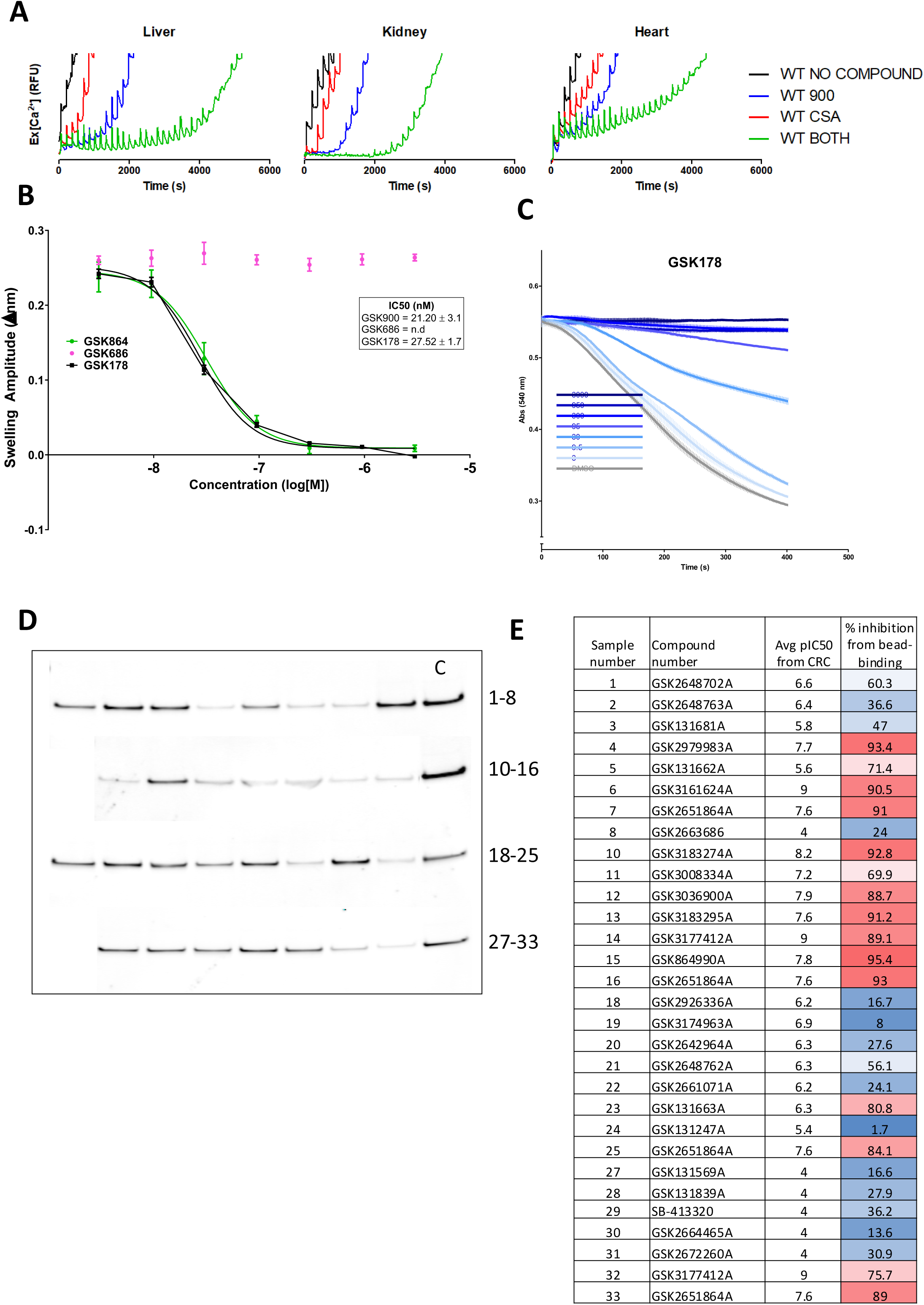
**A)** CRC of mouse liver, kidney or heart mitochondria at 2 nmol CaCl_2_ per mg protein per injection treated with 10 μM GSK900 (blue), CsA (red), GSK900+CsA (green) or DMSO (black). **B)** Dose response of GSK686 and GSK178 in swelling assay as in Figure 1D. **C)** Swelling assay of rat liver mitochondria with dose response of GSK178 as in Figure 1B. **D,E)**. GSK178 beads were incubated with HEK293 mitochondrial lysate in the presence of 30 μM compounds listed in **(E)** in a competition format, before amount of NLRX1 captured by GSK178 monitored by Western Blot **(D)**. % inhibition by each compound, noted by sample number in **(D)** and **(E)**, calculated by densitometry versus control (DMSO) in each row of samples.

**Supplementary Figure 3:**
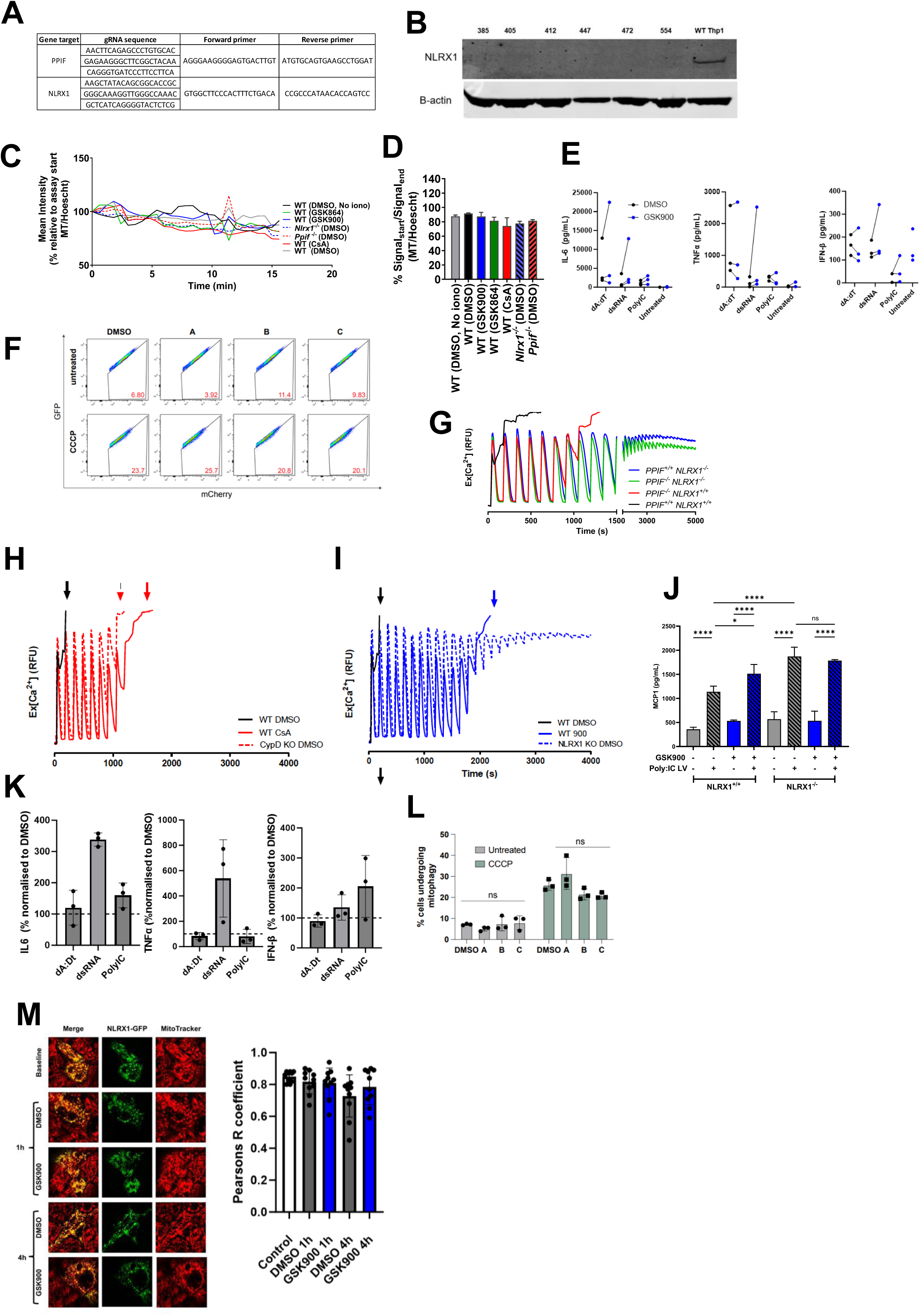
**A)** Guide and primer sequences used in CRISPR editing methods. **B)** Western blot confirmation of deletion of NLRX1 in THP-1 cells. Each clone noted by clone number. Cells were pooled for use. **C,D)** Mitotracker Red signal over time during mPTP cellular imaging assay in Figure 4B. Quantification of signal in **(D)**. **E)** Raw data for panel K; IL-6, TNFα and IFN-β as measured by Luminex from primary human macrophages pre-treated with 10 μM GSK900 before 24 h of immune stimuli as stated. **F)** Raw flow cytometry plots used to generate panel L, measuring fluorescence of mitophagy marker in transfected ARPE-19 cells treated with CCCP. **G)** High dose CRC assay of HEK293 isolated mitochondria from wildtype (black), NLRX1 KO (green), PPIF^-/-^ (red) or NLRX1^-/-^/PPIF^-/-^ (blue) cells. **H)** High dose CRC assay in mitochondria from wildtype HEK293 cells treated with vehicle (black) or CsA (red), and mitochondria from Ppif-/- HEK293 cells treated with DMSO (red, dashed). **I)** High dose CRC assay in wildtype mitochondria from HEK293 cells treated with vehicle (black) or GSK900 (blue), and mitochondria from NLRX1 KO HEK293 cells treated with DMSO (blue, dashed). **J**) MCP-1 release by wildtype or *NLRX1^-/-^* THP-1 cells after Poly:IC-LyoVec incubation for 24 h, with 10 μM GSK900 or DMSO. **K**) IL-6, TNFα and IFN-β release by primary human macrophages pre-incubated with DMSO or 10 μM GSK900 for 1 h, after 24 h of immune stimuli as indicated; raw data in panel E. **L**) CCCP-induced mitophagy in ARPE-19 cells expressing the mito-QC reporter, co-treated with vehicle, GSK900, GSK864 or GSK686 for 24 h; raw data in panel F. **M**) Wildtype HEK293 cells expressing NLRX1-GFP stained with Mitotracker Red at baseline, 1 or 4 h after vehicle or 10 μM GSK900; colocalisation as Pearson’s R (Coloc2); scale bar 10 μm. **(J)** where the statistics are relative to the vehicle condition (two-way ANOVA with Tukey’s multiple comparison test). Data in **(M)** are representative of at least 10 images, containing at least 5-10 cells per image. Each replicate in **(M)** is an average of all cells per image

**Supplementary Figure 4:**
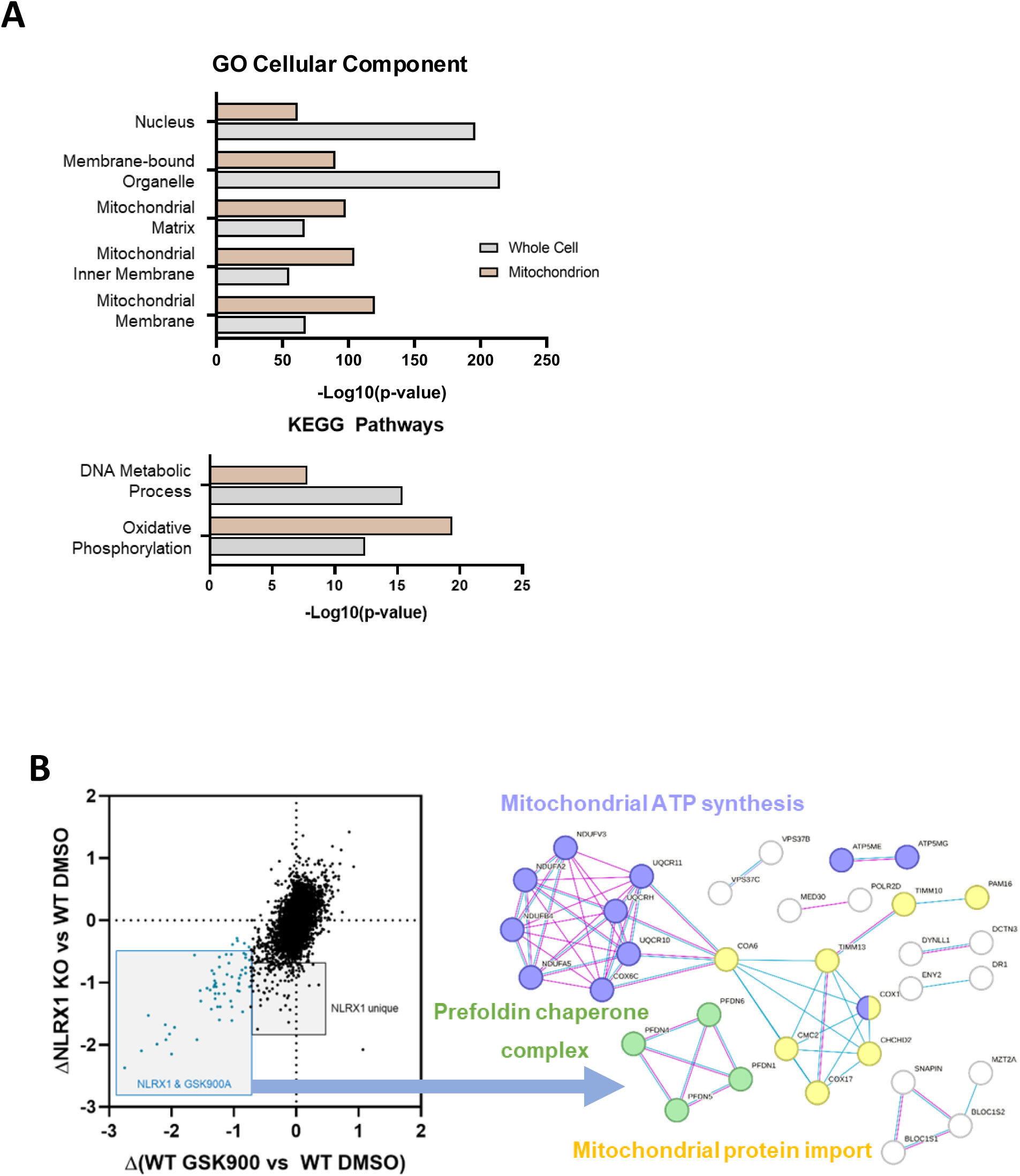
**A)** Cellular compartment gene ontology (GO) and KEGG pathway enrichments of whole cell and isolated mitochondrion samples. **B)** StringDB enrichment of correlated proteins exhibited in Figure 6.E. with associated GOs highlighted

**Supplementary Figure 5:**
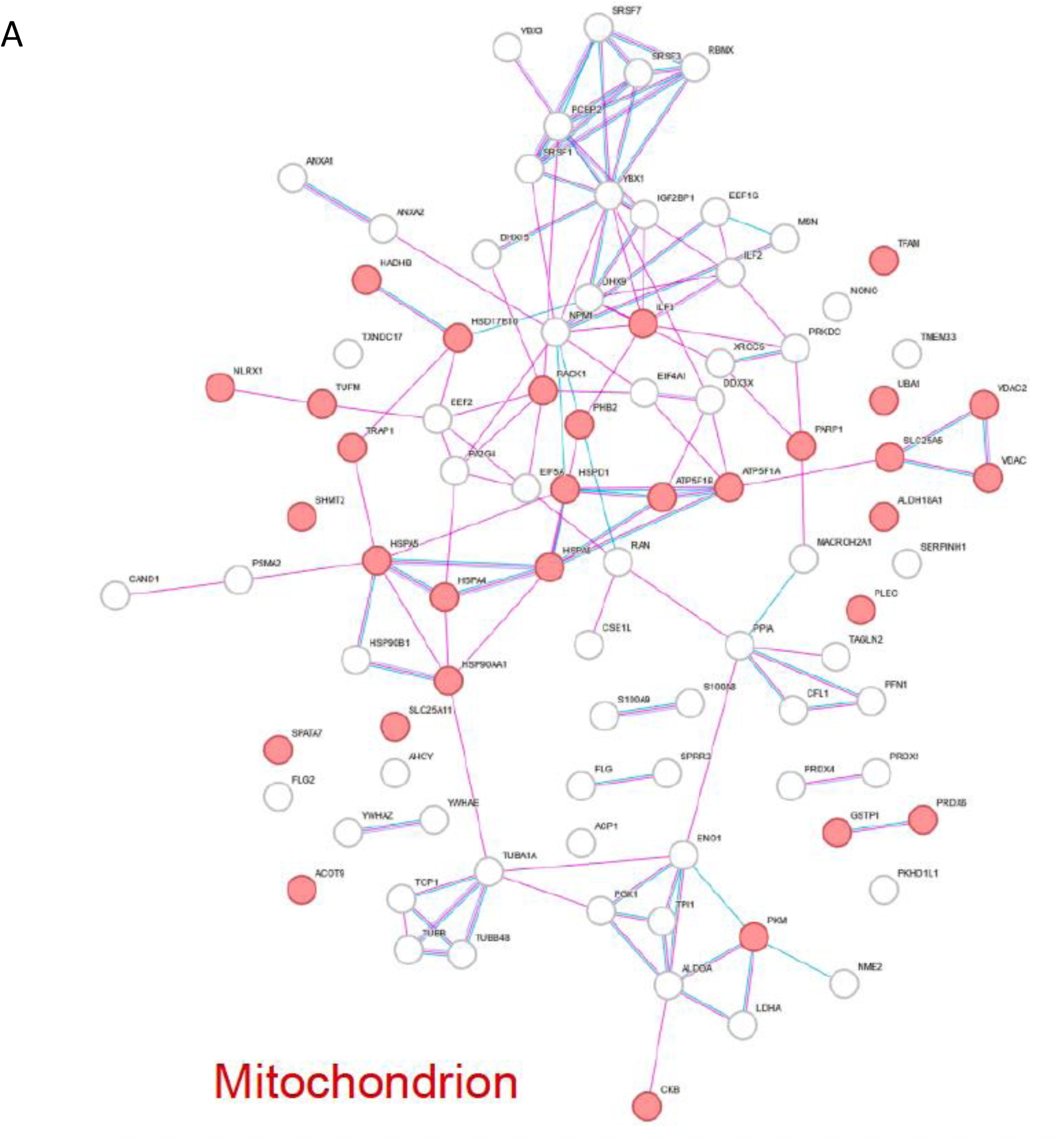
**A)** SDS-page gel separation of Co-IP samples and uncaptured fractions **B)** StringDB full interactome as detected by proteomics of captured fraction after co-immunoprecipitation of NLRX1 from wildtype HEK293 lysate.

**Supplementary Figure 6:**
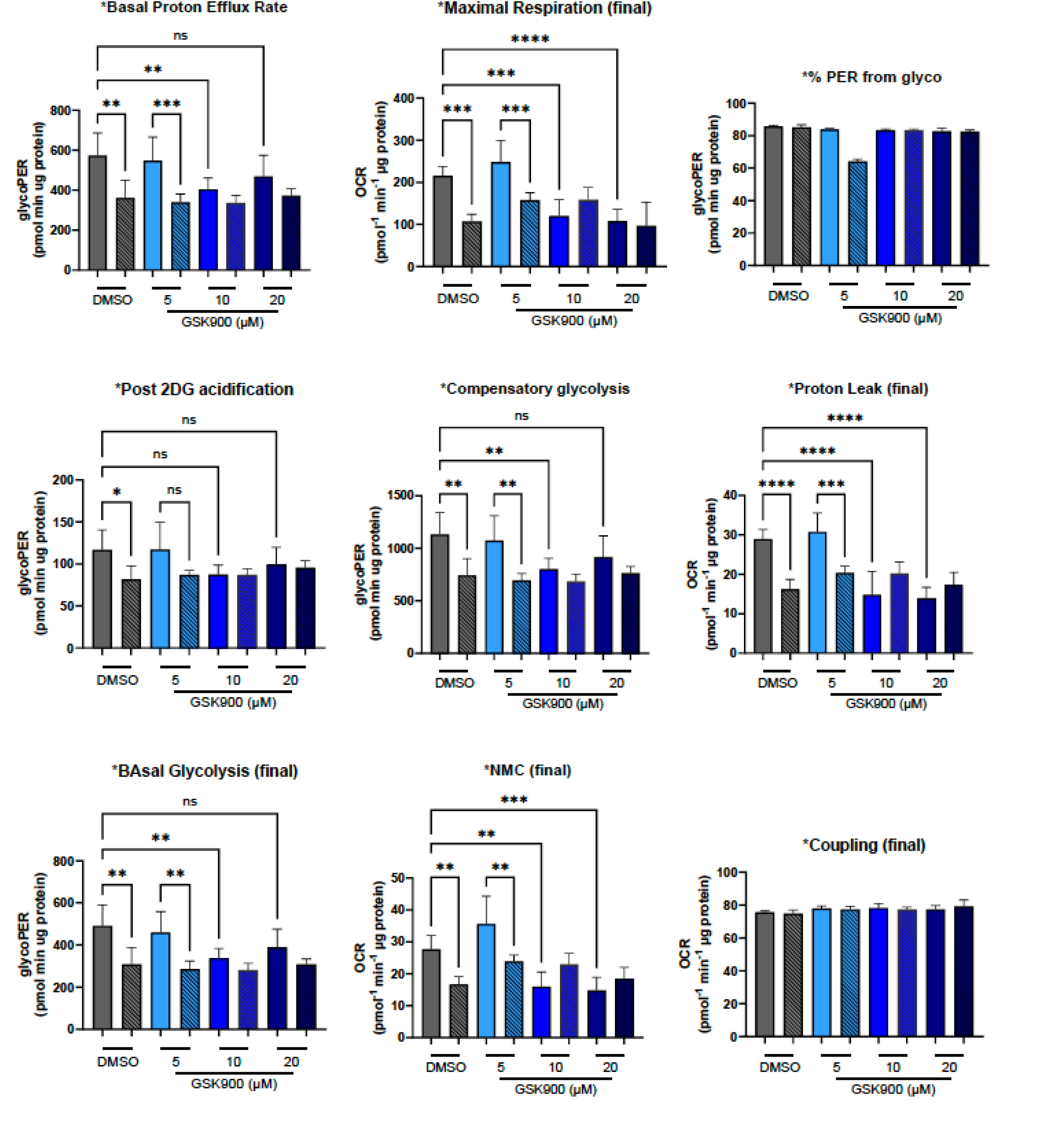
**A-J)** Seahorse parameters as measured by Wave software from Mitochondrial Stress Test and Glycolytic Rate test. All normalised to ug protein as measured by BCA assay. **A)** Basal proton efflux rate, **B)** Maximal respiration, **C)** % PER from glycolysis, **D)** post-2DG acidification, **E)** compensatory glycolysis, **F)** proton leak, **G)** Basal glycolysis, **H)** non-mitochondrial consumption, **I)** coupling efficiency (%). Wildtype cells are solid, *NLRX1^-/-^* are in dashed. Treatment with DMSO in grey, GSK900 in blue. * = *P*<0.05, ** = *P*<0.01, ***= *P*<0.001, **** = *P*<0.0001 as determined by a two-way ANOVA with Bonferroni multiple comparisons. Data is presented as mean ± S.D of at least three biological replicates

**Supplementary Figure 7:**
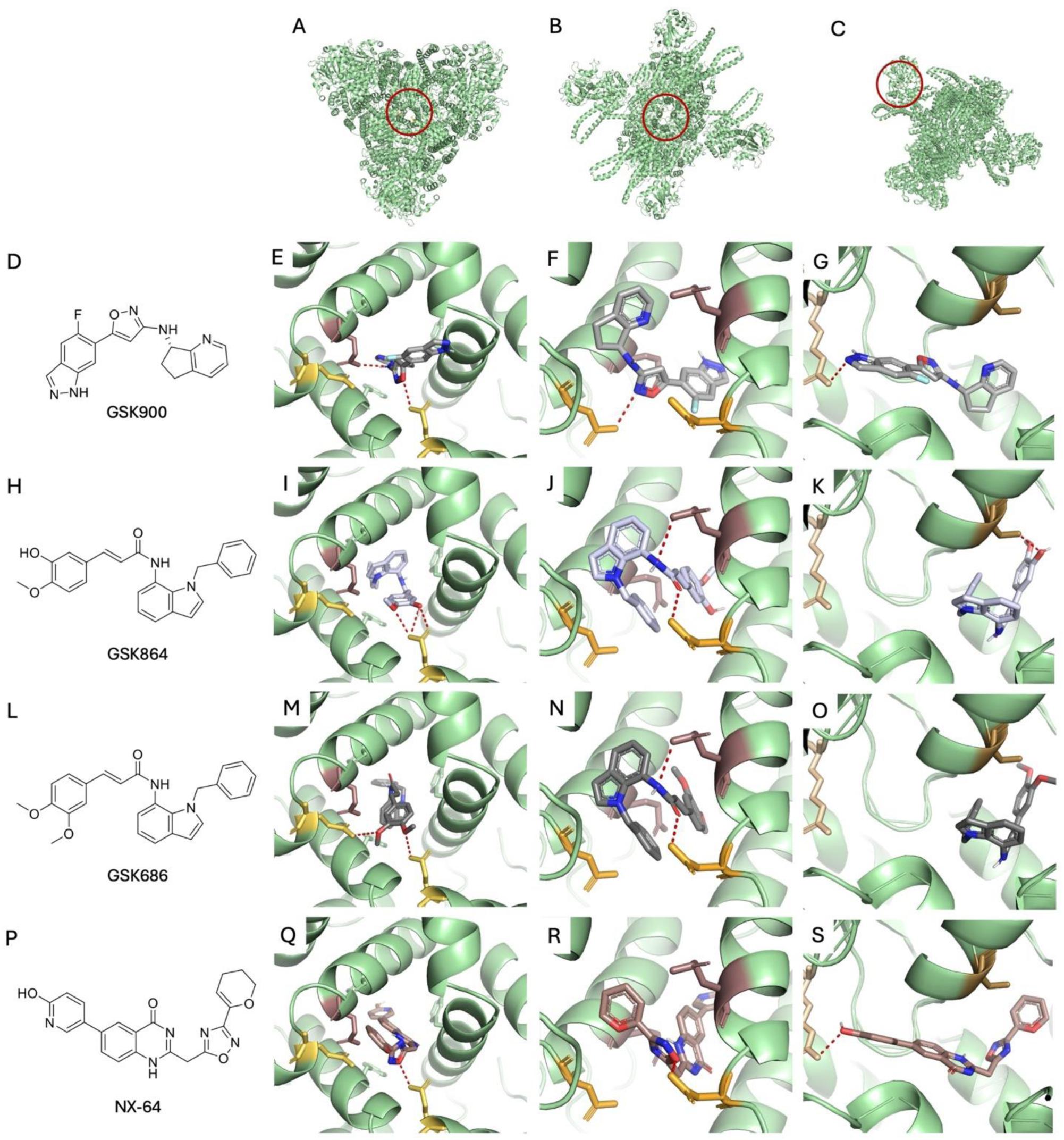
Binding interactions of small molecules with NLRX1. (A) Trimer interface where binding site is circled in red. (B) Dimer interface where binding site is circled in red. (C) ATP site at the N-terminal of NLRX1 where binding site is circled in red. (D) [name of the compound I]. (E) Interactions of [name] at the trimer interface of NLRX1 (−10.86 kcal/mol). (F) Interactions of [name] at the dimer interface of NLRX1 (−10.57 kcal/mol). (G) interactions of [name] at the ATP site of NLRX1 (−10.17 kcal/mol). (H) [name of the compound II]. (I) Interactions of [name] at the trimer interface of NLRX1 (−10.71 kcal/mol). (J) Interactions of [name] at the dimer interface of NLRX1 (−10.98 kcal/mol). (K) interactions of [name] at the ATP site of NLRX1 (−10.75 kcal/mol). (L) [name of the compound III]. (M) Interactions of [name] at the trimer interface of NLRX1 (−10.29 kcal/mol). (N) Interactions of [name] at the dimer interface of NLRX1 (− 10.99 kcal/mol). (O) interactions of [name] at the ATP site of NLRX1 (−9.809 kcal/mol). (P) NX-64-3 (Q) Interactions of NX-64-3 at the trimer interface of NLRX1 (−11.13 kcal/mol). (R) Interactions of NX-64-3 at the dimer interface of NLRX1 (−11.85 kcal/mol). (S) interactions of NX-64-3 at the ATP site of NLRX1 (- 11.38 kcal/mol).

